# Estrogen Drives Melanocortin Neurons To Increase Spontaneous Activity and Reduce Sedentary Behavior

**DOI:** 10.1101/794792

**Authors:** William C. Krause, Ruben Rodriguez, Bruno Gegenhuber, Navneet Matharu, Andreas N. Rodriguez, Adriana M. Padilla-Roger, Kenichi Toma, Candice B. Herber, Stephanie M. Correa, Xin Duan, Nadav Ahituv, Jessica Tollkuhn, Holly A. Ingraham

## Abstract

Estrogen depletion in rodents and humans leads to inactivity, unhealthy fat accumulation, and diabetes^1,2^, underscoring the conserved metabolic benefits of estrogen that inevitably decline with aging. In rodents, the preovulatory surge in 17β-estradiol (E2) temporarily allows energy expenditure to outpace energy intake, thus coordinating increased physical activity with peak sexual receptivity. To investigate how estrogen rebalances energy allocation in females, we examine estrogen receptor alpha (ERα) signaling in the ventrolateral ventromedial hypothalamic nucleus (VMHvl)^3–7^. We uncover a small population of VMHvl^ERα^ neurons expressing the melanocortin-4 receptor (MC4R) that integrates estrogen and melanocortin signals and projects to arousal centers in the hippocampus and hindbrain, enabling bursts of physical activity. ERα recruitment to the *Mc4r* gene promotes upregulation of *Mc4r* in VMHvl neurons during the preovulatory surge or following E2 treatment. We leveraged three models to stimulate VMHvl^MC4R^ neurons, restore MC4R signaling in the VMHvl of hyperphagic MC4R null females, or increase *Mc4r* levels in the VMHvl by CRISPR-mediated activation. All models increase spontaneous activity, whereas silencing VMHvl^MC4R^ neurons blunts normal activity. Direct activation of the VMHvl^MC4R^ node overrides the inactivity and hypometabolism following hormone depletion. These data extend the impact of MC4R signaling – the most common cause of monogenic human obesity^8^ – beyond the regulation of food intake. Our findings also rationalize reported sex differences in melanocortin signaling, including the greater disease severity of MC4R insufficiency in women^9^. The hormone-dependent node identified here illuminates the power of estrogen in motivating behavior during the female reproductive cycle and for maintaining an active lifestyle.

---

ERα signaling in VMHvl is a major determinant of energy usage in females, but the pathways and precise neuronal subsets that prioritize energy utilization over storage are unclear. To establish that maximal physical activity in females depends on ERα signaling in the VMHvl, we ablated ERα in the VMHvl or in the arcuate nucleus (ARC) of adult *Esr1^fl/fl^* female mice using stereotaxic delivery of AAV-Cre-GFP (VMHvl^ERαKO^, ARC^ERαKO^). Control female littermates received similarly targeted AAV-GFP injections (VMHvl^Control^ or ARC^Control^). Reduced ambulatory activity was observed in VMHvl^ERαKO^, but not ARC^ERαKO^, females during the dark (active) cycle that corresponded to a modest increase in body weight and a slight reduction in interscapular brown adipose tissue (iBAT) Ucp1 expression (Fig. 1a and Extended data 1.1c). While we previously showed that ARC^ERαKO^ females exhibit a surprisingly high bone mass phenotype4, no changes in daily food intake were noted in either VMHvl^ERαKO^ or ARC^ERαKO^ cohorts (Fig. 1a). Normal food consumption, particularly in ARC^ERαKO^ females, might suggest that extra-ARC sites^10^ mediate the anorexigenic effects of estrogen or that mouse strains/institutional housing conditions used here mask the expected hyperphagia in these females. When considered alongside other developmental ERα knockout models, these data demonstrate an ongoing requirement for ERα in the adult VMHvl to maximize daily patterns of physical activity in female mice.

**Fig. 1.**
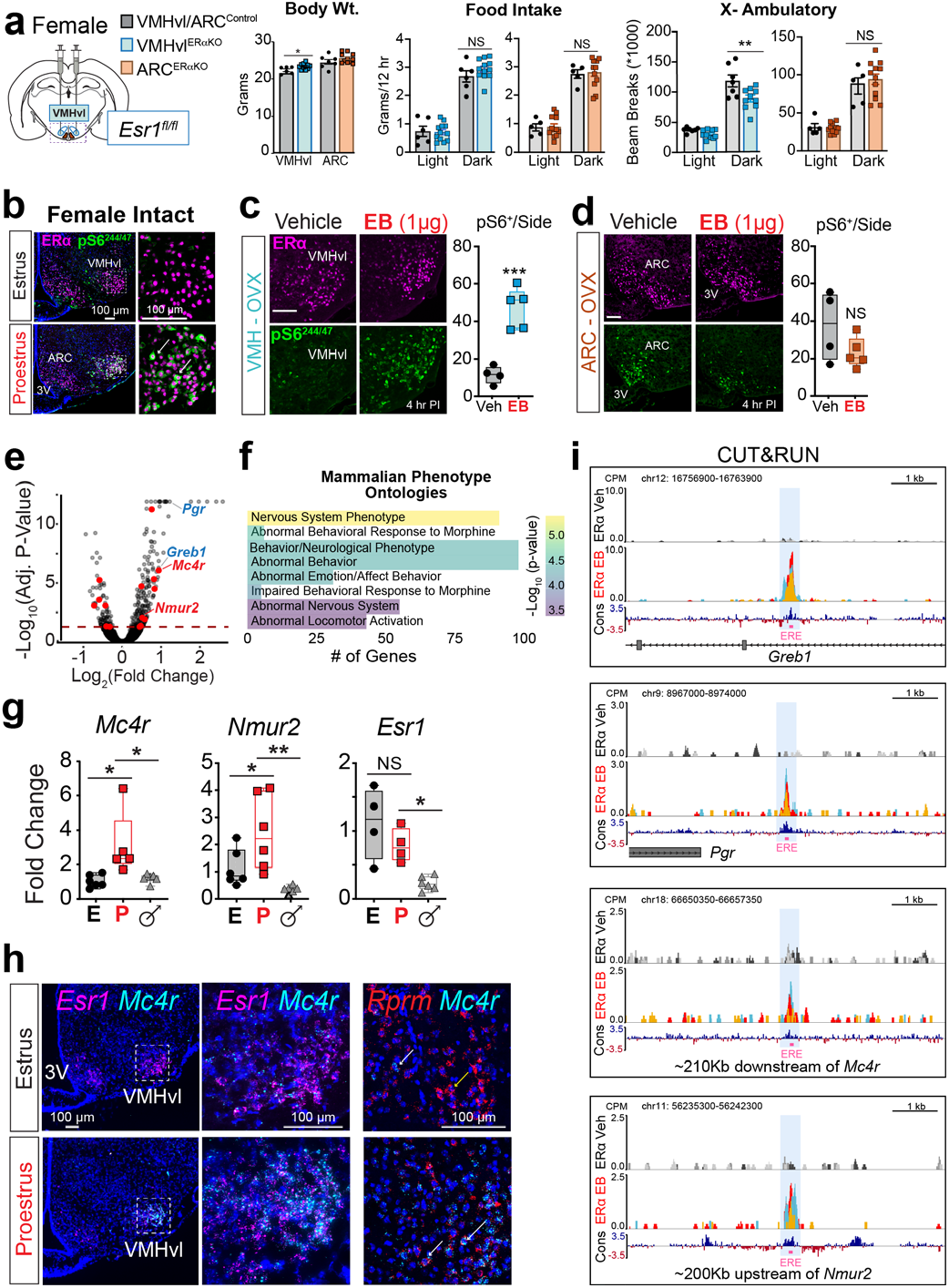
VMHvl neurons are sensitive to estrogen and maintain energy expenditure in adult females. **a,** Spatially-restricted ERα knockout and control mice were generated by stereotaxic delivery of AAV2-GFP or AAV2-Cre-GFP, respectively to the VMHvl or the ARC of *Esr1^fl/fl^* adult females. Quantification of body weight, food intake, and ambulatory activity in VMHvl^ERαKO^ (blue), ARC^ERαKO^ (orange), and their respective controls (grey). **b,** Immunofluorescent detection of ribosome phosphorylation (pS6^244/47^, green) and ERα (magenta) in the hypothalamus of intact females staged by vaginal cytology to be in estrus or proestrus. Representative images and quantification of pS6^244/47^ **c,** in the VMHvl and **d,** in the ARC of OVX female mice treated with vehicle or EB. **e,** Peptide ligand-binding receptors (red) are enriched among 287 EB-sensitive VMHvl genes (Benjamini-Hochburg adjusted P<0.05, dashed line). **f,** Mammalian Phenotype Ontology terms most significantly enriched among EB-sensitive genes. **g,** qPCR analysis of the indicated target genes in VMHvl from estrus females (E), proestrus females (P), and males (♂). **h,** Fluorescent ISH for *Mc4r* (cyan) *Esr1* (magenta) in VMHvl of estrus and proestrus females. Boxed areas are shown in right panels. Separate sections showing ISH for *Mc4r* and *Rprm* (red) with arrows indicating representative cells expressing both (white) or only Rprm (yellow). **i,** CUT&RUN CPM-normalized coverage track showing EB-specific ERα binding sites containing EREs (pink boxes) within the *Greb1* locus (3/3 replicates), *Pgr* locus (3/3 replicates), *Nmur2* locus (2/3 replicates), and *Mc4r* locus (2/3 replicates) (MACS2, q < 0.01) in 400,000 sub-cortical brain nuclei collected from vehicle and EB (5 μg) treated gonadectomized mice. *P<0.05; **P<0.01; ***P<0.001.

Hormone responsiveness of VMHvl^ERα^ neurons was visualized across the estrous cycle by monitoring phosphorylated ribosomal protein S6 (pS6) during estrus (low E2) and proestrus (high E2). pS6 signals increased substantially in the VMHvl during proestrus or following a single estradiol benzoate (EB) injection into ovariectomized (OVX) females (Fig. 1b, c), but were negligible in females during estrus, in females lacking ERα, or in intact males without exogenous EB (Extended data 1.2a,b), underscoring a complete dependence of this pS6 response on both E2 and ERα. Induction of pS6 by EB in VMHvl^ERα^ neurons occurred slowly (>2 hrs post-hormone injection), suggesting a classical genomic mechanism. No changes in pS6 induction were detected in adjacent ARC^ERα^ neurons (Fig. 1c), demonstrating that VMHvl^ERα^ neurons are highly, and selectively sensitive to E2. That pS6 signals are induced in the VMHvl during the E2 surge but suppressed when energy supplies decline (e.g., during fasting or leptin deficiency)^11^, argues that estrogen recalibrates the perceived energetic state, setting the stage for a change in physical activity.

Candidate downstream mediators of E2 signaling were identified after profiling the VMHvl transcriptome in OVX mice treated with vehicle or EB (Fig. 1e). Among the 287 differentially expressed genes (DEGs), we noted enrichment of those involving peptidergic G-protein coupled receptor signaling, as well as behavioral phenotypes, including altered locomotor activity (MP:0003313, adjusted *P*= 3.19E-4), (Fig. 1f and Extended data 1.3a). In particular, EB affected the expression of several metabolically relevant neuropeptide receptors, including *Mc4r*, *Nmur2*, *Npy1r*, and *Ghsr*, and known estrogen and/or sex-dependent genes (*Greb1*, *Pgr*^4^). Both MC4R^12,13^ and NMUR2^14^, affect locomotor behavior, and interestingly, *Mc4r* and *Nmur2* expression was induced during proestrus (P) but near absent during estrus (E) and in intact males (Fig. 1g). Given that MC4R coordinates the balance between energy storage and allocation, as well as previously observed sex differences in *Mc4r* knockouts and loss-of-function mutations in both mice^13,15^ and humans^9,16^, we hypothesized that melanocortin signaling could underly the control of physical activity by VMHvl^ERα^ neurons. We confirmed that *Mc4r* expression increases during proestrus or in response to EB and was colocalized with VMHvl neurons expressing ERα (*Esr1*) or *Rprm*, a VMHvl female-specific marker that is non-estrogen responsive^7^ (Fig. 1h and Extended data 1.3b-d).

We further established that *Mc4r* is a direct transcriptional target of ERα using CUT&RUN (Cleavage Under Targets and Release Using Nuclease), which detects transient in vivo binding events within heterogenous tissues17. As expected, hormone-dependent ERα-chromatin interactions were detected in *Greb1* and *Pgr*, two genes that are strongly expressed throughout hypothalamic and forebrain structures that control sex-dependent behaviors^18^. Despite the exceedingly small number of estrogen-responsive neurons that coexpress *Mc4r*, the high sensitivity afforded by CUT&RUN enabled detection of two conserved ERα binding sites within the *Mc4r* locus. The first site is located −210kb downstream of the transcript with a complete estrogen response element (ERE) consensus sequence. The second site resides in the proximal promoter and is composed of an ERE half-site (+84 bp) and a recognition site for the trans-acting transcription factor 1 (Sp1, +67 bp, Fig. 1i, and Extended Data 1.4a,b); these two binding motifs are known to coordinate estrogen-dependent global regulation of ERα target genes^19^. A full ERE consensus was detected +200kb upstream of *Nmur2*, consistent with upregulation of *Nmur2* during proestrus (Fig. 1i and Extended Data 1.4b). Together, these data establish a direct molecular link between estrogen and MC4R signaling and indicate that estradiol dynamically regulates neuropeptidergic input to VMHvl^ERα^ neurons.

We then compared ERα and MC4R co-expression in the VMHvl and other candidate brain regions using a Cre-dependent fluorescent reporter, Ai14, to label neurons under the control of Mc4r-t2a-Cre (Ai14^*Mc4r*^)^20^. VMHvl^ERα/MC4R^ neurons represent ~40% of the broader VMHvl^ERα^ population (Fig. 2a). In contrast to the nearly perfect overlap in the VMHvl, concordance of Ai14^*Mc4r*^ and ERα in the medial amygdala (MeA) is lower, and the degree of Mc4r induction during proestrus appears weaker (Extended data 2.1a). No overlap could be discerned in two other regions of interest – the paraventricular hypothalamus (PVH), a primary site that couples MC4R with its effects on food intake that expresses little-to-no ERα, and the lateral hypothalamic area (LHA) that harbors a small subset of LHA^MC3R/MC4R^ neurons linked to increased locomotor activity^21^.

**Figure 2.**
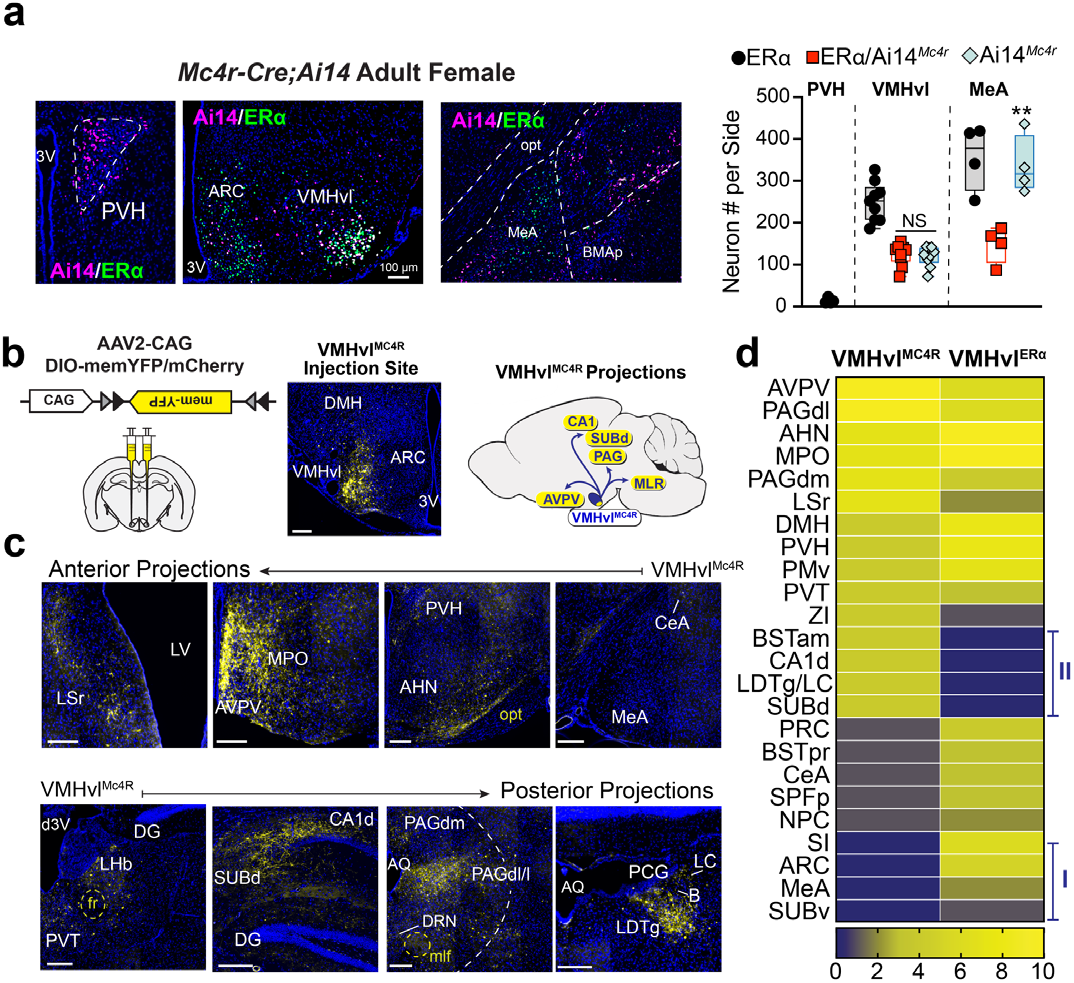
VMHvl^MC4R^ neurons are a moleculary and anaomically distinct subset of VMHvl^ERα^ neurons. **a,** Representative coronal brain images of Ai14^*Mc4r*^ female mice stained for ERα (green) and Ai14 (magenta) in the PVH, VMHvl and MeA and quantification of neurons labeled by Ai14^*Mc4r*^ alone or co-expressing ERα/ Ai14^*Mc4r*^. **b,** Overview of Cre-dependent labeling strategy to map VMHvl^MC4R^ neuron projections (left) and example of targeting of viral vector to VMHvl (middle) with major projections depicted (right). **c,** Labeled projections from the VMHvl to anterior (upper row) and posterior (lower row) neuroanatomical regions; scale bars = 200 μm. Images are representative of bilateral targeting of VMHvl (n = 5). **d,** Semi-quantitative comparison of projection intensities from VMHvl^MC4R^ neurons with those previously identified for VMHvl^ERα^ neurons^22^ reveals neuroanatomical targets that are VMHvl^ERα^-specific (Cluster I) or VMHvl^MC4R^-specific (Cluster II). Anatomical abbreviations listed in Extended data Table 1. **P<0.01.

Given that VMHvl^MC4R^ neurons represent a subset of the VMHvl^ERα^ population, we asked if afferent projections—as labeled by Cre-dependent, membrane-targeted YFP (mYFP) (Fig. 2b)—are distinct from the larger VMHvl^ERα^ population. Overall, we identified many (~84%) of the same major ascending and descending projections reported for VMHvl^ERα^ neurons^22^, but failed to see strong targeting to the ARC and medial amygdala (MeA) (Fig. 2c,d Cluster I and Extended data 2.1b). Unexpectedly, VMHvl^MC4R^ neurons projected to the dorsal CA1 (CA1d) and the adjacent subiculum (SUBd) (Fig. 2c,d Cluster II), a hippocampal region controlling locomotor speed in mice^23^ and containing “speed cells” whose firing rate correlates with velocity^24^. As expected, strong projections from VMHvl^MC4R^ neurons to the pre-motor periaqueductal grey (PAG) region were detected, but interestingly showed an innervation pattern largely restricted to the lateral and dorsolateral PAG columns (PAGdl/l) that are associated with escape behaviors^25^, while conspicuously avoiding the ventrolateral PAG (PAGvl), a midbrain region associated with freezing and defensive behaviors^26^ (Fig. 2d and Extended data 2.1d). In addition, VMHvl^MC4R^ neurons projected to the pontine central gray, a hindbrain region containing a cluster of nuclei that mediate sexual receptivity, locomotor arousal, and wakefulness^27–29^. Our findings establish VMHvl^MC4R^ neurons as an overlapping but distinct subset of the VMHvl^ERα^ population projecting to discrete brain regions that are functionally consistent with spontaneous locomotion in response to fluctuating hormone levels.

While Mc4r expression has been reported in the VMH^20, 30, 31^, little is known about its role in the ventrolateral subregion. As VMHvl^ERα^ neurons promote energy utilization via physical activity^3^ and thermogenesis^6^, and also modulate glucose homeostasis^32^, we artificially activated these VMHvl^MC4R^ neurons using Designer Receptors Exclusively Activated by Designer Drugs (DREADDs, AAV-DIO-hM3Dq-mCherry) to assess their functional output. Cre-dependent DREADDs were injected bilaterally in the VMHvl of *Mc4r-t2a-Cre*-positive and Cre-negative littermates (Fig. 3a). Administration of the DREADD ligand, clozapine-n-oxide (CNO), during the normally inactive lights on period increased spontaneous physical activity in both female and male VMHvl^MC4R::hM3Dq^ mice but not in VMHvl^Cre-^ controls (Fig. 3b,c, and Extended data 3.1a). The response to a single injection of CNO in VMHvl^MC4R::hM3Dq^ mice lasted approximately five hours, and during this time distance traveled by both sexes jumped more than 700% while sedentary behavior dropped significantly (Extended Data 3.1b).

**Figure 3.**
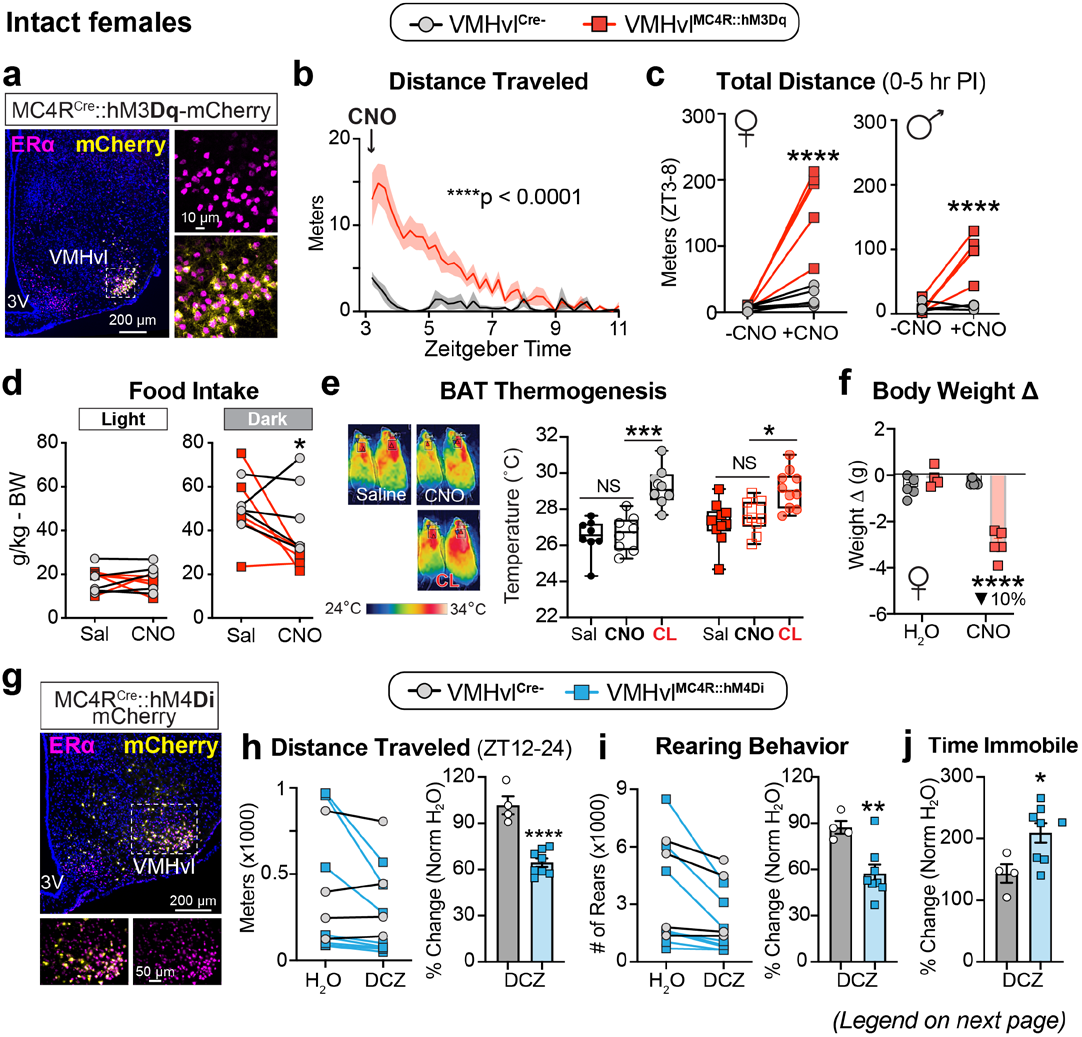
Stimulating VMHvl^MC4R^ neurons promotes energy expenditure by increasing spontaneous physical activity. **a,** ERα and mCherry expression in the VMHvl following targeted injection of Cre-dependent AAV-hM3Dq-mCherry into a female *Mc4r-t2a-Cre* mouse. **b,** Distance traveled over time in intact female VMHvl^MC4R::hM3Dq^ and VMHvl^Cre-^ mice post-injection (PI) of CNO. **c,** Total distance traveled over ZT3-ZT8 for female and male mice without (-CNO) or 5 hours post-injection of CNO (+CNO). **d,** Five hour food consumption in female mice, normalized to body weight, following saline (Sal) or CNO injection during early light (ZT4) and beginning of dark (ZT12) periods. **e,** Thermal imaging of BAT surface temperatures for VMHvl^Cre-^ (left mouse) and VMHvl^MC4R:: hM3Dq^ (right mouse) females with graph showing BAT surface responses 30 and 45 min after injection of Sal, CNO, or CL-316,243. **f,** Weight change in intact female mice following 24 hour administration of plain drinking water (H^2^O) or CNO-laden water (CNO). **g,** ERα and mCherry expression in the VMHvl following targeted injection of Cre-dependent AAV-hM4Di-mCherry into a female *Mc4r-t2a-Cre* mouse. **h-j,** Distance traveled (h), number of rearing episodes (i), and change in immobile time (j) during the dark period in intact female mice following administration of plain or DCZ-laden drinking water. *P<0.05; **P<0.01; ***P<0.001; ****P<0.0001.

Aside from the change in movement, other metabolic functions were insensitive to activation of the VMHvl^MC4R^ neurons. A modest effect on food intake was observed only during the active period (lights off) (Fig 3d). In contrast to the β-3 adrenergic agonist, CL-316-243, CNO did not increase iBAT temperature or iBAT expression of *Ucp1* mRNA in either VMHvl^MC4R::hM3Dq^ or VMHvl^Cre-^ controls (Fig. 3,e and Extended data 3.1c). VMHvl^MC4R::hM3Dq^ mice also responded equivalently to a glucose tolerance test (GTT, i.p.) when treated concurrently with either saline or CNO, although the higher body weights inherent to the *Mc4r-t2a-Cre* line lowered their glucose tolerance (Extended data 3.1d,e). Providing CNO in the drinking water over a 24 hour period led to a nearly 10% drop in body weight in VMHvl^MC4R::hM3Dq^ females with a corresponding increase in activity (Fig. 3f, Supplementary Video, and Extended data 3.2a-c). CNO treatment extended over an eight day period resulted in a 13% drop in body weight in VMHvl^MC4R::hM3Dq^ females (Extended data 3.2d). Collectively, these findings show that VMHvl^ERα/MC4R^ neurons constitute a potent node to specifically promote physical activity, which can be artificially engaged in both sexes. Conversely, targeting the VMHvl with inhibitory DREADDs (AAV-DIO-hM4Di-mCherry) (Fig. 3g) reduced spontaneous physical activity and increased sedentary behavior during the dark period in VMHvl^MC4R::hM4Di^ mice administered DREADD ligand (Fig. 3h-j). Thus, consistent with the reduction in ambulatory activity observed in VMHvl^ERαKO^ females, VMHvl^MC4R^ neuron activity is essential for generating normal nighttime activity levels in female mice.

Declining estrogen levels during aging are associated with lowered energy expenditure, increased sedentary behavior, and metabolic decline. We therefore asked if DREADD-activation of VMHvl^MC4R^ neurons would override the deleterious consequences of estrogen depletion. Chemogenetic activation of VMHvl^MC4R^ neurons in OVX females fully restored physical activity parameters and promoted significant weight loss over a short 24 hr period (Fig. 4a-c and Extended data 4.1a,b). As the combined impact of estrogen depletion and overnutrition in women can impair glucose homeostasis^1^, we challenged OVX VMHvl^MC4R::hM3Dq^ females with high fat diet (HFD) for 5 wks (Extended data 4.1c). Whereas fasting glucose and insulin tolerance were unchanged following CNO treatment in OVX females on standard chow diet (Extended data 4.1d), both parameters improved notably after a single bout of CNO-induced activity in HFD-fed VMHvl^MC4R::hM4Di^ females (Fig. 4d and Extended data 4.1e). Further, chronic stimulation of VMHvl^MC4R^ neurons in obese, sedentary OVX females resulted in a rapid, dramatic weight loss (Fig. 4e and Extended data 4.1f), accompanied by fasting blood glucose, a drop in cellular adiposity of gonadal fat, and reduced plasma cholesterol (Fig. 4f, g and Extended data 4.1g, h), all markers of improved metabolic health. Food intake was unaffected (Fig. 4h). Taken together, engagement of the VMHvl^MC4R^ activity node reduces body weight in OVX females, independent of diet, and improves metabolic health in the face of a dietary challenge and estrogen depletion.

**Fig. 4.**
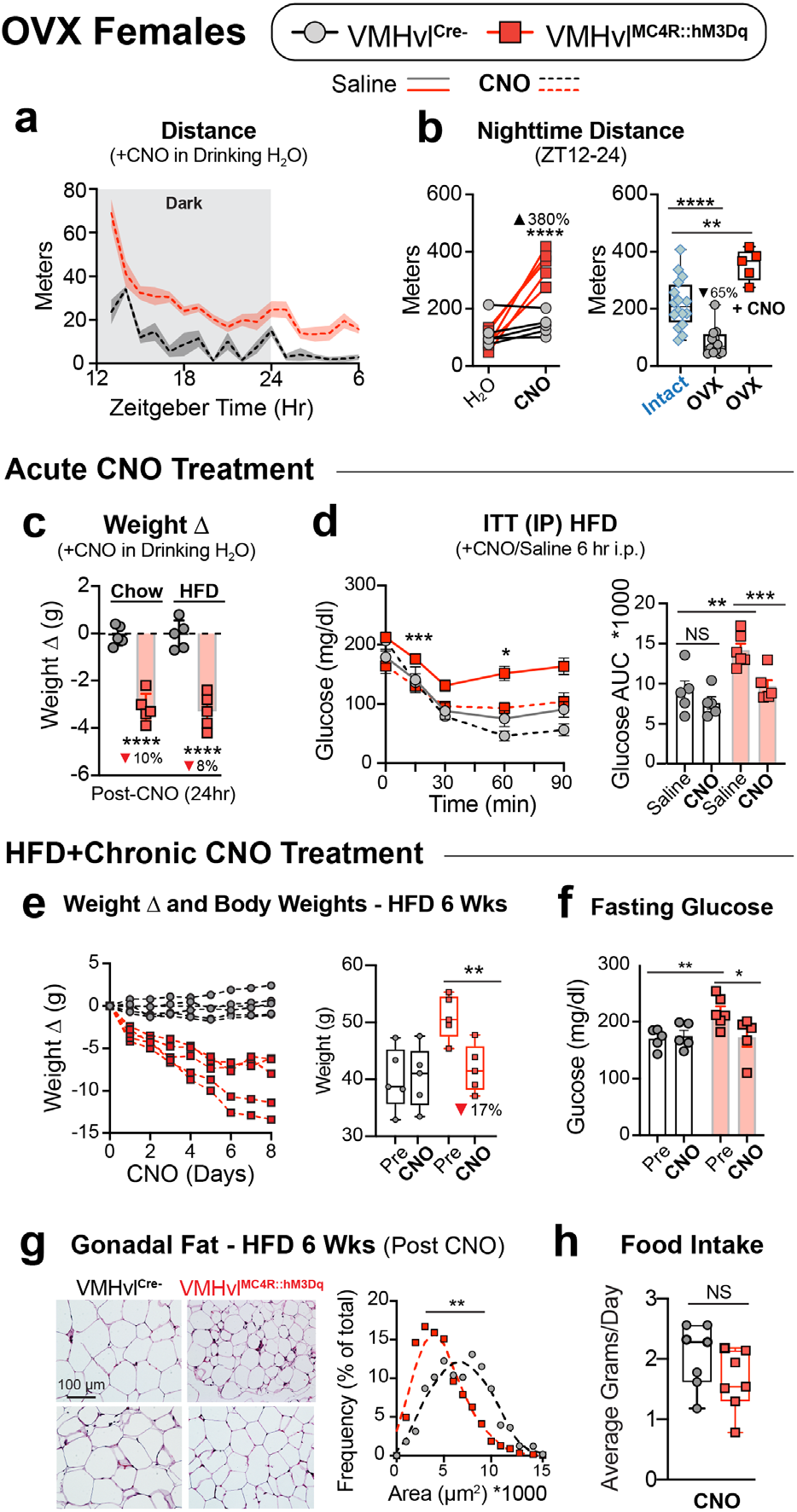
Activation of VMHvlMC4R neurons in OVX females restores physical activity levels and reverses markers of metabolic dysfunction following high-fat diet feeding. **a,** Distance traveled over time in OVX female VMHvl^Cre-^ and VMHvl^MC4R::hM3Dq^ mice during administration of plain H_2_O or CNO-H_2_O. **b,** Cumulative distance during dark period in OVX females following CNO treatment (left panel), and then compared to intact (blue) and OVX (grey) female mice (right panel). **c,** Body weight change in OVX females fed standard diet (Chow) or high-fat diet (HFD) following 24-hour administration of CNO in H_2_O. **d,** Blood glucose (left) and AUC (right) values during 90 min ITT test on HFD-fed OVX females performed 6 hours post fasting and post injection with saline or CNO. **e,** Body weight change during 8-day chronic CNO treatment of OVX/HFD mice (left) with average body weights plotted on Day 0 (Pre) and on Day 8 (Post) (right). **f,** Six hour fasting blood glucose levels in OVX/HFD mice before and after 8 days of chronic CNO. **g,** Representative H&E staining of gonadal white adipose from OVX/ HFD mice following 8 days of CNO in H_2_O with graph showing frequency distribution of adipocyte area. **h,** Average daily food intake per mouse of group-housed, HFD-fed OVX females during chronic CNO administration with each point representing a separate daily measurement. *P<0.05; **P<0.01; ***P<0.001;****P<0.0001.

To determine whether melanocortin signaling itself activates this VMHvl activity node, we first examined its responsiveness to MT-II, a synthetic MC4R agonist, in female mice pretreated with EB or vehicle for five hours. VMHvl cFOS expression was highest when a single, high dose of MT-II (400 μg, i.p.) was preceded by EB, establishing that MC4R modulates VMHvl neuronal activity (Fig. 5a,b). As expected, MT-II also increased cFOS-positive cells in the PVH. We next used the Cre-dependent *Mc4r^loxTB^* allele^33^ in combination with the *Sf1-Cre* transgene, which is expressed in the VMH but does not overlap with *Mc4r* expression in the periphery (Extended data 5.1a) to restore Mc4r specifically in the VMH of otherwise null mice (*Mc4r^Sf1-Cre^*). In response to EB, this Cre-dependent rescue approach increased *Mc4r* expression in VMHvl^*Esr1*^ neurons, similar to wild type females (*Mc4r^+/+^*) (Fig. 5c). At weaning, body weights were equivalent between groups (Extended data 5.1b). Subsequently, rescued *Mc4r^Sf1-Cre^* and null *Mc4r^loxTB^* adult females exhibited phenotypes attributed to the PVH, including severe hyperphagia and increased body length33 (Fig. 5d and Extended data 5.1c). Restoring *Mc4r* to the VMHvl attenuated both the overt obesity and sedentary behavior in female but not male MC4R null mice (Fig. 5e-g). Consistent with estrogen-dependent expression, these data support the hypothesis that the female VMHvl relies on MC4R signaling to promote spontaneous activity.

**Fig. 5.**
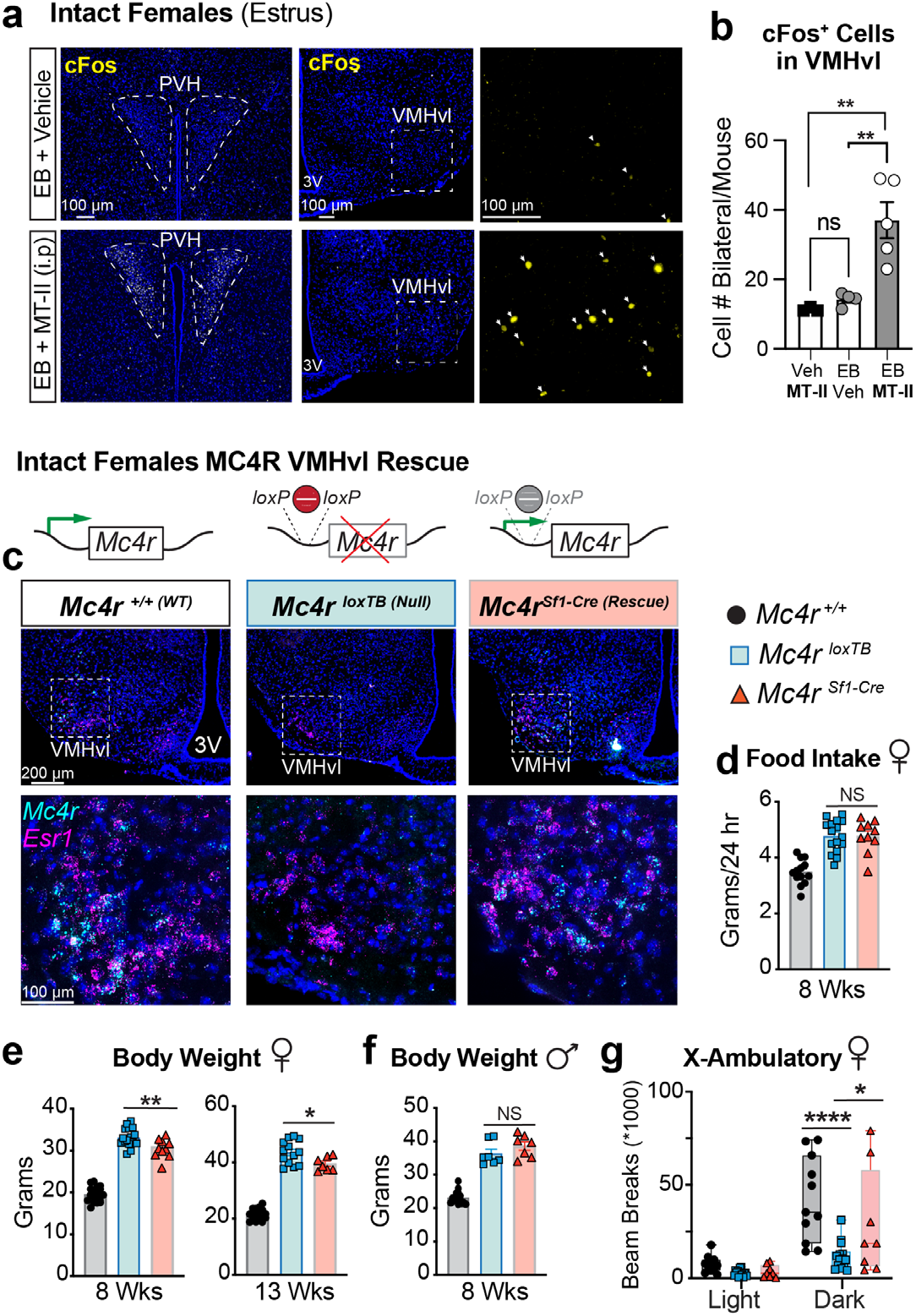
EB-treated VMHvl neurons are sensitive to melano-cortin and restoration of VMHvl^MC4R^ signaling is sufficient to increase spontaneous physical activity and decrease obesity specifically in female mice. **a,** Representative cFos immunostaining (arrows) in the PVH and VMHvl of female mice treated with EB or vehicle -/+MT-II. **b,** Quantification of VMHvl cFos induction. **c,** Fluorescent ISH comparing *Esr1* and *Mc4r* expression patterns in EB-treated *Mc4r^+/+^*, *Mc4r^loxTB^*, and *Mc4r^Sf1-Cre^* females. **d,** Food intake in *Mc4r^+/+^*, *Mc4r^loxTB^*, and *Mc4r^Sf1-Cre^* female mice. **e,** Body weights in 8- and 13-week-old females. **f,** Body weights in 8-week-old *Mc4r^+/+^*, *Mc4r^loxTB^*, and *Mc4r^Sf1-Cre^* males. **g,** Light and dark period ambulatory movement in adult *Mc4r^+/+^*, *Mc4r^loxTB^*, and *Mc4r^Sf1-Cre^* females. *P<0.05; **P<0.01; ***P<0.001; ****P<0.0001.

To verify that MC4R signaling is an integral component of the hormone-responsive VMHvl activity node, CRISPR-mediated activation (CRISPRa) was employed to increase Mc4r expression in the VMHvl. Previously, Mc4r CRISPRa targeting to the PVH was shown to normalized gene dosage and restore energy balance in haploinsufficient *Mc4r^+/-^* mice^34^. Here, wild type female and male mice were stereotaxically injected with a dual vector system containing the Mc4r promoter guide RNA targeting the ERE half-site (AAV-Mc4r-Pr-sgRNA) and an endonuclease-deficient dCas9 tethered to the VP64 transcriptional activator (AAV-dCas9-VP64) to selectively upregulate *Mc4r* expression in the VMHvl (Fig. 6a). Control mice received dCas9-VP64 but not the sgRNA. Delivery of dual Mc4r-CRISPRa-viral vectors to the VMHvl was confirmed post-mortem by mCherry signal and *Mc4r* ISH or by qPCR quantification that resulted in a moderate, long-lived induction of *Mc4r* in both sexes (Fig. 6b and Extended data 6.1a,b). CRISPRa^*Mc4r*^ females traveled on average two-fold more in the dark phase compared to control females, with increased movement persisting for at least 17 weeks post-injection (Fig. 6c). CRISPRa^*Mc4r*^ was unable to restore normal activity in OVX females (Extended data 6.1f,g). Activity in CRISPRa^*Mc4r*^ males also increased (Fig. 6d). Increased physical activity in CRISPRa^*Mc4r*^ mice was restricted to the dark, thus preserving the normal diurnal activity pattern. In accordance with elevated levels of spontaneous activity during the dark phase, CRISPRa^*Mc4r*^ mice spent less time immobile during this period (Extended data 6.1c). Although body weights (Fig. 6e) and BAT activity (Extended data 6.1d) were equivalent, a modest but significant increase in daily food intake was detected in female CRISPRa^*Mc4r*^ mice (Fig. 6f). After weeks of elevated daily activity and increased mechanical loading in CRISPRa^*Mc4r*^ females, cortical bone thickness and bone volume were higher (Fig. 6g and Extended data 6.1e). Thus, sidestepping ERα and directly increasing *Mc4r* dosage in the VMHvl permanently increases spontaneous activity behavior in both sexes.

**Fig. 6.**
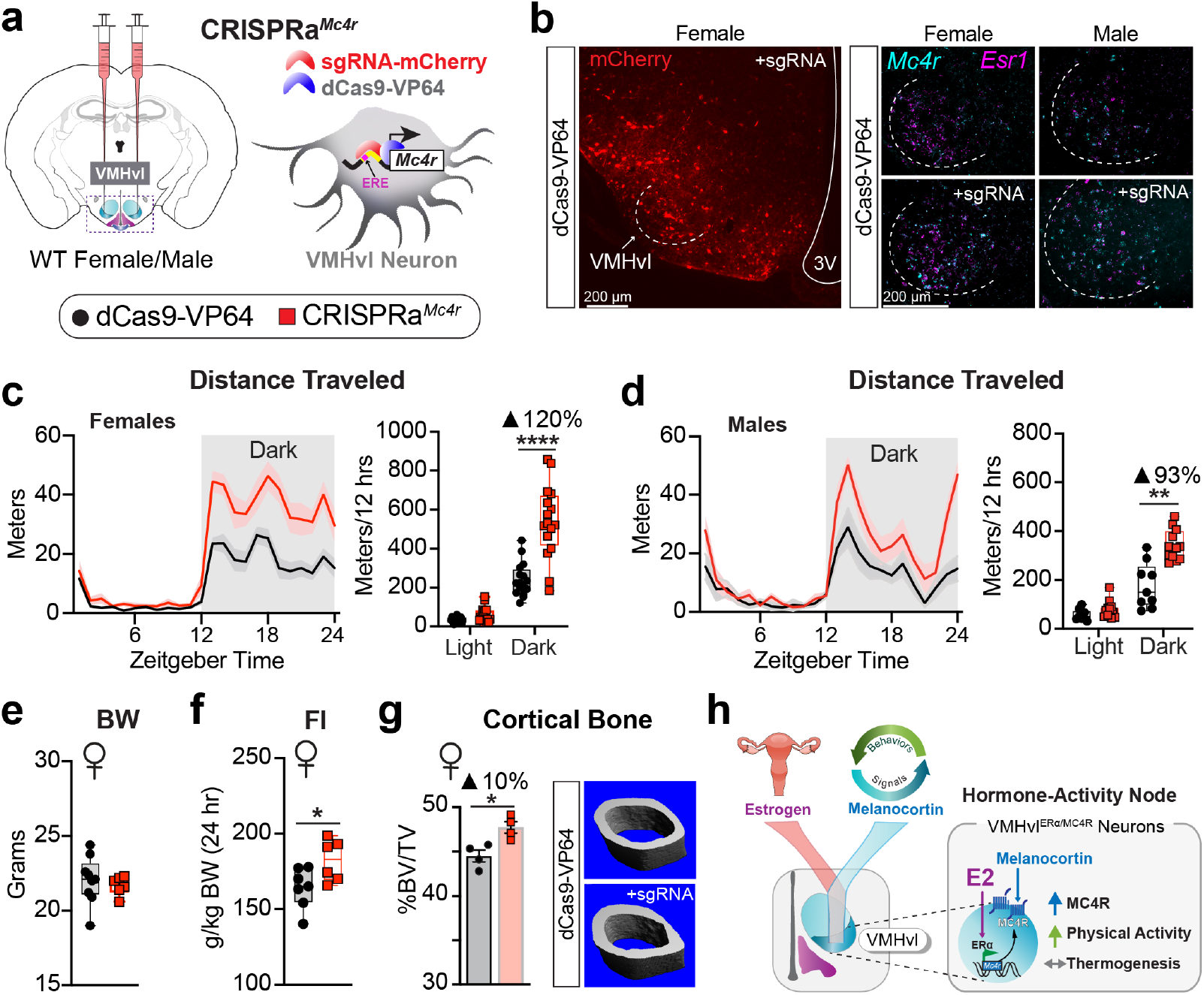
Bypassing E2-dependent *Mc4r* induction by CRISPRa chronically increases physical activity levels in both male and female mice. **a,** Schematic overview of injection and dual vector CRISPRa^*Mc4r*^ system targeting the *Mc4r* promotor region containing the ERE half-site. Control mice received dCas9-VP64 alone. **b,** mCherry fluorescence and *Mc4r*/*Esr1* fluorescent ISH to confirm VMHvl delivery of sgRNA and spatially restricted target gene induction, respectively. **c-d,** Distance traveled for female and male CRISPRa^*Mc4r*^ (n=6 female, 3 male) and control (n=5 female, 4 male) mice in home cages over 24 hours with total distance traveled in the light and dark phases for three most active runs from each mouse. **e,** Body weights (BW) of CRISPRa^*Mc4r*^ and control female cohorts fed standard chow diet ad lib. **f,** Average daily food intake (FI) normalized to body weight in female cohorts measured over a 4-day period. **g,** Cortical bone volume fraction for female cohorts 4 months post-infection. **h,** Upregulation of *Mc4r* by ERα in response to E2 in VMHvl^ERα/MC4R^ neurons integrates estrogen and melanocortin signaling to generate a specialized hormone-dependent activity node in females. *P<0.05; **P<0.01; ***P<0.001; ****P<0.0001.

## DISCUSSION

Here we identify an estrogen-sensitive VMHvlERα/ MC4R node that maximizes daily patterns of spontaneous physical activity in female mice. Within this hormone-responsive node, we show that MC4R is a crucial intermediary component coupling estrogen and energy expenditure through its direct transcriptional regulation by ERα (Fig. 6h). Thus, as *Mc4r* expression increases during the preovulatory period (high E2), sensitivity to melanocortin rises in the VMHvl resulting in the well-documented estrogen-dependent spike in activity observed in rodents^35^ and ruminants^36^. Responses in VMHvl^MC4R^ neurons would be amplified further by the positive actions of estrogen on production of alpha melanocortin stimulating hormone (αMSH) in the ARC, the endogenous MC4R agonist^37^. Collectively, our data underscore the potent role of estrogen to promote activity during a critical point in the reproductive cycle.

Although correct Mc4r dosage is known to be critical for normal function and metabolic regulation in both humans and rodent models, endogenous signals that increase its expression have not been described. Our data identify estrogen as a potent signal that increases *Mc4r* expression, akin to the gain-of-function human MC4R variants that attenuate receptor internalization and protect against weight gain^38^. The high degree of conservation in the consensus ERE and ERE/Sp1 binding motifs observed in the mammalian *Mc4r* locus suggests that estrogen similarly upregulates MC4R expression in humans. Furthermore, our findings are entirely consistent with the hormone-dependency of MC4R synthetic agonists to stimulate sexual behaviors in estrogen-primed female rodents^39^ and enhance libido in premenopausal women suffering from hypoactive sexual desire disorder^40^. Conversely, estrogen depletion would render VMHvl neurons unresponsive to melanocortins by limiting MC4R expression. Such melanocortin resistance might contribute to the increased sedentary lifestyle associated with menopause^41^, while acknowledging that the benefits of estrogen for a healthy metabolism likely involve additional MC4R-independent processes^42^. Our data reconcile noted sex differences in the MC4R literature. For example, women harboring MC4R variants exhibit a higher risk for metabolic disease^9^. Experiments in mice indicate a female-specific requirement for MC4R independent of the PVH that point to missing MC4R regulatory sites for optimizing energy homeostasis in females^13,33^. That we counteract the severe obesity in Mc4r-/-females after restoring MC4R to the VMHvl argues strongly that a critical missing component of melanocortin signaling for females is the VMHvl^MC4R^ node.

Once engaged, these specialized VMHvl^MC4R^ neurons target CNS regions important for female reproductive behavior, such as the AVPV, PMv, and PAG. However, identification of VMHvlMC4R projections to sites that regulate the speed of locomotion, orientation, and arousal including the hippocampal area encompassing the SUBd and CA1d^23,24^, and areas of the hindbrain associated with arousal and motor output^27–29^ expand greatly the possible functional outputs of the VMHvl. Whether these VMHvl outputs contribute to psychiatric disorders in women that commonly emerge at life stages involving changes in reproductive hormones (e.g. postpartum depression and premenstrual dysphoric disorder) remains to be determined.

Despite the pronounced increase in physical activity observed in CRISPRa^*Mc4r*^ females, body weights remained stubbornly unchanged in the face of small increases in daily food intake. On the other hand, DREADD activation of VMHvl^MC4R^ neurons rapidly reduced body weight in estrogen-depleted OVX females and improved metabolic parameters; however, the rate of weight loss is not sustainable (Extended data 5.1f). Our findings reinforce the notion that engagement of adaptive responses often limits the effectiveness of exercise on weight loss^43,44^. Nonetheless, decreasing sedentary behavior, independently of body weight, reduces the risk of several metabolic- and age-related co-morbidities, including heart disease, frailty, cancer, and infectious diseases^45^. As such, the extremely durable increase in spontaneous physical activity achieved by the non-transgenic CRISPRa^*Mc4r*^ approach provides a unique preclinical model to explore the motivational aspects and health benefits of an active lifestyle. The central mechanism identified here illustrates how estrogen signaling leverages MC4R signaling to drive bursts of physical activity during peak female sexual receptivity. Our findings underscore the benefits of estrogen in minimizing sedentary behavior and provoke further discussion about hormone replacement therapies in postmenopausal women.

## METHODS

### Mice

All experiments were conducted in accordance with UCSF IACUC guidelines and the approved protocol for the Ingraham Lab. *Mc4r^loxTB^* mice and the *Ai14^fl/fl^* reporter mice were purchased from Jackson Laboratories and maintained on a C57BL/6J background. *Mc4r-t2a-Cre* mice were a generous gift from B. Lowell (BIDMC) and were maintained on a C57BL/6J background. *Esr1^fl/fl^* were maintained on a mixed background, and *Sf1-Cre*^46^ mice were maintained on a C57BL/6N in the lab as previously described^3,4^. Wild type mice used for CRISPRa studies were on a pure C57BL/6J background. For *Mc4r* rescue experiments, the *Sf1-Cre* was contributed through female mice. Mice were housed on a 12:12 hour light cycle (lights on 6:00; lights off 18:00) and had ad libitum access to standard chow (LabDiet #5058) or high-fat diet (Research Diets #D12492). CUT&RUN experiments were performed on adult male (8-12 weeks of age) gonadectomized C57Bl6/J wild type mice obtained from Jackson Laboratory. Three weeks post-gonadectomy, animals were injected subcutaneously with either corn oil (vehicle) or 5 μg of estradiol benzoate and sacrificed after 4 hours. For each biological replicate, brain dissections were pooled from 5 animals.

#### Stereotaxic Injections

AAV2-Cre-GFP and AAV2-GFP were purchased from the UNC Vector Core (Chapel Hill, NC). AAV2-hM3Dq-mCherry and AAV2-hM4Di-mCherry were gifts from Bryan Roth and viral preparations were purchased from Addgene (viral prep # 44361-AAV2; http://n2t.net/addgene:44361; RRID:Addgene_44361 and viral prep # 44362-AAV2; http://n2t.net/addgene:44362; RRID:Addgene_44362; Addgene, Watertown, MA)^47^. For axonal tracing, AAV2-CAGs-FLEX-membrane-YFP-WPRE.hGH was re-engineered to fuse YFP with a C-terminal farnesylation tag to enhance the membrane labeling^48^. AAV2 was prepared using a standard polyethylene glycol gradient followed by cesium chloride density gradient centrifugation protocol^49^ to reach a titer of (1×10E13 GC/ML). AAVdj-dCas9-VP64 and AAVdj-Prm-Mc4r-sgRNA, were generated by the Stanford Gene Vector and Virus Core and details of vector constructs are as previously described ^34^. Adult mice were secured in a Model 1900 stereotaxic frame (David Kopff Instruments, Tujunga, CA), and 250-600 nL of virus was injected bilaterally at the following coordinates: For the VMHvl - A-P: Bregma −1.48 mm, M-L: Bregma +/−0.85 mm, D-V: skull −5.9 mm. For the ARC: A-P: Bregma −1.58 mm, M-L: Bregma +/−0.25 mm, D-V: skull −5.8 mm.

For all surgeries regardless of viral vectors used, mice recovered for at least two weeks prior to any metabolic or behavioral assays. For projection labeling, mice were allowed to express the reporter for 5-8 weeks before tissue collection. At the conclusion of the experiments, mice were euthanized and the brains were collected to confirm proper targeting. Any mice absent of correctly targeted fluorescent protein expression were excluded from subsequent analyses. Water-soluble CNO (HB6149; Hello Bio, Princeton, NJ) was administered by IP injection (0.3 mg/kg in sterile saline) or in the drinking water (0.25 mg/mL). CNO-laden drinking water was replaced every 48 hours. Water-soluble DCZ (HB9126; Hello Bio, Princeton, NJ) was administered in the drinking water (0.1 mg/mL) ^50^.

### Estrous Cycle Staging and EB Treatment

Reproductive stages in female mice were determined by comparing relative amounts of leukocytes, epithelial cells and cornified epithelial cells collected by vaginal lavage. Stage assessments were made daily between ZT3 and ZT5. Brains from estrus or proestrus females were collected between ZT7 and ZT10 and processed for immunofluorescence, ISH, or qPCR.

Adult female mice (>8 week old) were OVX. Estradiol benzoate (Cayman Chemical, 10006487) was dissolved in DMSO and diluted in sesame oil (Sigma, S3547). Mice received a subcutaneous injection of either 1ug EB in 150 μL sesame oil or 150 μL of sesame oil with an equivalent amount of DMSO. Control mice received a subcutaneous injection of 150 μL of sesame oil with an equivalent amount of DMSO. To minimize changes in VMH gene expression or signal transduction associated with fear/anxiety, mice were handled daily in a manner that simulated injection for at least 5 days prior to EB/Vehicle treatment and tissue collection. For cFOS analyses, mice were treated with 400μg MT-II (Bachem) by i.p. injection, and brains were collected 1-1.5 hours later.

### RNA-seq and qPCR

Brains from OVX females treated with EB (n = 4) o vehicle (n = 3) were rapidly dissected into ice-cold PBS with 0.1% DEPC. Coronal brain sections (250 μm thick) were cut on a vibratome and transferred to glass slides so that the VMH could be visualized and manually microdissected. Isolated tissue was flash frozen and stored at −80°C. RNA was prepared using the RNeasy Micro kit (Qiagen). Sequencing libraries were constructed using the TRIO RNA-seq Library Preparation kit (TECAN) using 15 ng of input RNA. Equal amounts of each sample library were multiplexed and sequenced (50 bp single-end reads) on a single flow cell lane HiSeq 4000 (Illumiina). Demultiplexed reads were aligned to the mouse genome (mm10) using HISAT251 and counted using HTSeq52. Finally, differential gene expression testing was performed using DESeq2^53^.

Isolated RNA, prepared as described above, was converted to cDNA using the SuperScript III reverse transcriptase (Invitrogen). Target genes were amplified using specific primers (Mc4r forward: 5-GCCAGGGTACCAACATGAAG-3 and reverse: 5-ATGAAGCACACGCAGTATGG-3; Nmur2 forward: 5-CCTCCTTCCTCTTCTACATCCT-3 and reverse: 5-AGTCACTTTGTCTGCCTCAA-3; Esr1 forward: 5-GAACGAGCCCAGCGCCTACG-3 and reverse: 5-TCTCGGCCATTCTGGCGTCG-3; and Ucp1 forward: 5-CACGGGGACCTACAATGCTT-3 and reverse: 5-TAGGGGTCGTCCCTTTCCAA-3). Ct values were normalized to cyclophilin (Ppib, forward primer: 5-TGGAGAGCACCAAGACAGACA-3 and reverse primer: 5-TGCCGGAGTCGACAATGAT-3), and relative expression levels were quantified using the comparative CT method. Individual values, representing the VMHvl or iBAT from 1 mouse are the average of 2 technical replicates.

### CUT&RUN

ERα CUT&RUN was performed on 400,000 nuclei isolated from BNSTp, POA, and MeA tissue via density gradient centrifugation54. Briefly, tissue was homogenized 15x with a loose pestle in a glass homogenizer containing Homogenization Medium (250 mM sucrose, 25 mM KCl, 5 mM MgCl2, 20 mM Tricine-KOH, 1 mM DTT, 0.15 mM spermine, 0.5 mM spermidine, 1X Roche EDTA-free protease inhibitor cocktail, pH 7.8). 0.3% IGEPAL CA-630 was added, and the tissue was further dounced 5x with a tight pestle. After douncing, the homogenate was filtered through a 40 μm strainer and mixed 1:1 with 50% OptiPrep solution (Millipore Sigma) prepared in Dilution Buffer (150 mM KCl, 30 mM MgCl2, 120 mM Tricine-KOH, pH 7.8). The homogenate was underlaid with 5 ml of 30% and 40% OptiPrep solution, respectively, and centrifuged at 10,000xg for 18 min at 4°C in an ultracentrifuge. ~2 ml of nuclei solution were removed from the 30 - 40% OptiPrep interface by direct tube puncture. Following nuclei isolation, 0.4% IGEPAL CA-630 was added to improve binding to concanavalin A magnetic beads (Bangs Laboratories BP531). CUT&RUN was performed on brain nuclei, according to the standard protocol17. Nuclei were washed twice in Wash Buffer (20 mM HEPES, pH 7.5, 150 mM NaCl, 0.1% BSA, 0.5 mM spermidine, 1X PIC) and incubated overnight on a nutator with ERa antibody (Millipore Sigma 06-935), diluted 1:100 in Antibody Buffer (Wash Buffer containing 2 mM EDTA). Nuclei were washed twice in Wash Buffer, and ~700 ng/ml protein A-MNase (pA-MNase) was added. After 1 hr incubation on a nutator at 4oC, the nuclei were washed twice in Wash Buffer and placed in a metal heat block on ice. pA-MNase digestion was initiated by 2 mM CaCl2. After 90 min, pA-MNase activity was stopped by mixing 1:1 with 2X Stop Buffer (340 mM NaCl, 20 mM EDTA, 4 mM EGTA, 50 μg/ml RNase A, 50 μg/ ml glycogen). Digested fragments were released by incubating at 37°C for 10 min, followed by centrifuging at 16,000xg for 5 min at 4°C. DNA was purified from the supernatant by phenol-chloroform extraction.

### CUT&RUN Library Preparation

CUT&RUN libraries were prepared using the SMARTer ThruPLEX DNA-seq Kit (Takara Bio), with the following PCR conditions: 72oC for 3 min, 85oC for 2 min, 98oC for 2 min, (98oC for 20 sec, 67oC for 20 sec, 72oC for 30 sec) x 4 cycles, (98oC for 20 sec, 72oC for 15 sec) × 10 cycles. Samples were size-selected with AMPure XP beads (1.5X right-sided and 0.5X left-sided) to remove residual adapter dimers and large DNA fragments. Individually barcoded libraries were multiplexed and sequenced with paired-end 75 bp reads on an Illumina NextSeq, using the High Output Kit.

### CUT&RUN Data Processing

Paired-end reads were trimmed with cutadapt55 to remove low-quality base-calls (-q 30) and adapters. Trimmed reads were aligned to mm10 using Bowtie256 with the following flags: --dovetail --very-sensitive-local --no-unal --no-mixed --no-discordant --phred33. After alignment, duplicate reads were removed using Picard (http://broadinstitute.github.io/picard/) MarkDuplicates (REMOVE_DUPLICATES=true). De-duplicated reads were filtered by mapping quality (MAPQ > 40) using samtools^57^ and fragment length (< 120 bp) using deepTools alignmentSieve^58^. After filtering, peaks were called using MACS2 callpeak^59^ with a q-value threshold of 0.01 and min-length set to 25. Individual replicate BAM files were normalized by counts per million (CPM) and converted to bigwig tracks, using deepTools bamCoverage (-bs 1, --normalize using CPM). CPM-normalized bigwig tracks for individual EB and vehicle samples (n=3 per condition) were plotted using Gviz^60^.

### In Situ Hybridization

For colorimetric ISH, antisense *Mc4r* probes were PCR amplified (forward primer: 5-ACTCTGGGTGTCATAAGCCTGT-3 and reverse primer: 5-TCTGTCCCCCACTTAATACCTG-3) from hypothalamic cDNA libraries, and in vitro transcribed with incorporation of digoxigenin-UTP (Roche) using the T7 or SP6 Riboprobe kit (Promega). 20 μm sections from fixed tissue were labeled and detected as previously described3. Fluorescent ISH was performed using RNAScope (ACD, Multiplex Fluorescent V2) according to the manufacturer’s protocol using the following probes: *Esr1* (478201), *Mc4r* (319181-C2), and *Rprm* (466071).

### Immunofluorescent Staining and Histology

Fixed CNS tissue was cryosectioned (20 μm) and stained overnight with primary antibodies against: ERα (EMD Millipore, #C1355 polyclonal rabbit or Abcam, #93021 monoclonal mouse), phospho-Serine 244/247 RPS6 (Invitrogen, #44-923G polyclonal rabbit), cFOS (Santa Cruz, SC-52 polyclonal rabbit), or red fluorescent protein (Rockland, 600-401-379 polyclonal rabbit). For detection, sections were labeled with species-appropriate secondary Alexa Fluor-coupled antibodies (Invitrogen, #A11029 and #A11037).

Fixed gonadal white adipose tissue was paraffin embedded, sectioned (5 μm), and stained with hematoxylin and eosin (H&E) by the Gladstone Histology and Light Microscopy core. Brightfield images were thresholded to define adipocyte borders, and the adipocyte area was quantified using ImageJ.

### Micro-Computed Tomography

Following perfusion fixation, femurs from CRISPRaMc4r and Control female mice were isolated. Volumetric bone density and bone volume were measured by μCT as previously described4.

### Metabolic and Activity Monitoring

Indirect calorimetry and food intake were measured in CLAMS chambers (Comprehensive Laboratory Animal Monitoring System, Columbus Instruments). Any spilled food that was not consumed was accounted for at the conclusion of the four day period spent in CLAMS.

Ambulatory activity in mice subjected to chemogenetic or CRISPRa manipulations was recorded via IR cameras and quantified using the ANY-maze behavioral tracking system (Stoelting). Prior to any measurements, mice acclimated to single-housing in the ANY-maze chambers for at least three days. For CRISPRa studies, activity tracking was continuously monitored for at least five 24hr periods.

Interscapular skin temperatures were measured using a FLIR-E4 handheld infrared camera (FLIR Systems, Inc. Wilsonville, Oregon) as previously described. Female mice were lightly anesthetized in groups of four or five in an anesthesia induction chamber and images were captured at baseline, 30 minutes, and 60 minutes post saline, CNO (0.3 mg/kg) or CL-316,243 (3 mg/kg) intraperitoneal injection.

Blood glucose and lipid assays were performed following a 6 hour fast (starting ~ZT2) during which mice were housed in clean cages with ad libitum water access. For glucose and insulin tolerance tests, fasted mice were injected with glucose (I.P., 1 g/kg) or inulin (I.P., 1 U/kg), respectively. Tail-blood samples were collected at baseline and every 15 minutes after glucose/insulin injection. Blood glucose levels were quantified using a hand-held glucometer (Roche, Accu-Check Compact). For triglyceride and cholesterol measurements, plasma was isolated from tail-blood and measured (3 μL in technical duplicates) using the Cholesterol Quantitation Kit (Sigma, MAK043) or the Triglyceride Quantification Colorimetric/Fluorometric Kit (Sigma, MAK266).

### Statistics

Statistical tests, excluding RNA-Seq analyses, were performed using Prism 8 (Graphpad). A description of the test and results are provided in Extended data Table 2. 1-way, 2-way, and repeated measures (RM) ANOVA multiple comparisons were performed using the Holm-Sidak post hoc test. Unless otherwise noted, data are presented as mean ± SEM.

## ACKNOWLEDGMENTS

We thank Dr. C. Paillart and T. McMahon for exceptional technical assistance with the running and data acquisition for the CLAMS and Any-maze systems. Additionally, we thank all members of the Ingraham Lab for their many insightful comments and discussions, Dr. O. Yabut for expertise in imaging, as well as Dr. C. Vaisse for insights on MC4R signaling. This work was supported by funding to H.A.I. (R01 DK121657, R01AG062331, GCRLE Senior Scholar Award 0320, UCSF Women’s Reproductive Health RAP Award, and NRSA NDSP P30-DK097748), to W.C.K. (American Heart Association Postdoctoral Fellowship 16POST27260361), to R.R. (IRACDA K12 GM081266 Program), to B.G. (2T32GM065094, F31MH124365), to N.M. (UCSF Mary Ann Koda-Kimble Innovation Seed Award, UCSF Catalyst Program), to X.D. (R01 EY030138), Research to Prevent Blindness and Klingenstein-Simons Neuroscience Fellowship to S.M.C (K01 DK098320, UCLA Women’s Health Center, UL1TR001881), to C.B.H (F32 DK107115-01A1, AHA Postdoctoral Fellowship 16POST29870011) to N.A. (R01 CA197139, R01 MH109907), to J.T. (R01 MH113628, SFARI600568). We wish to acknowledge the mouse metabolic core funded by P30 DK098722-01.

## AUTHOR CONTRIBUTIONS

W.C.K. designed experiments, analyzed data, and wrote the manuscript. R.R. performed thermal and glucose homeostasis analyses in mice. B.G. optimized, performed, and analyzed the CUT&RUN method for ERα binding in neurons. N.M. provided CRISPRa viral vectors and expert advice. A.R. performed histology and quantification of expression data. A.M.P-R. aided with chemogenetic data acquisition and analyses. C.B.H. analyzed bone and plasma lipid data. S.M.C. designed experiments, provided animal models, and analyzed data. K.T. and X.D. provided the AAV-DIO-mYFP vector. N.A. provided key unpublished reagents related to CRISPRa constructs and helped guide studies. J.T. optimized CUT&RUN method for ERα binding in neurons, performed analyses, and wrote the manuscript. H.A.I designed experiments, analyzed data, and wrote the manuscript.

