## Supplementary Figures and Tables for "Estrogen Drives Melanocortin Neurons To Increase Spontaneous Activity and Reduce Sedentary Behavior"

### Supplementary Information

**Supplementary Video:** CNO administered in the drinking water stimulates VMHvl$^{MC4R}$ neurons to increase physical activity.

**Extended Data Figure 1.1:** VMHvl pS6 induction depends on ERα and can be elicited in males by EB treatment.

**Extended Data Figure 1.2:** VMHvl$^{ER\alpha KO}$ affects brown adipose thermogenic gene expression in mice maintained at ambient temperature.

**Extended Data Figure 1.3:** EB-dependent regulation of *Mc4r*.

**Extended Data Figure 1.4:** ERα binding sites in EB-sensitive target genes contain conserved ERE consensus sequences.

**Extended Data Figure 2.1:** Sensitivity of *Mc4r* expression in the VMHvl and VMHvl$^{MC4R}$ neuron output.

**Extended Data Figure 3.1:** Chemogenetic activation of VMHvl$^{MC4R}$ neurons reduces sedentary behavior during the normally inactive light period but does not affect glucose homeostasis.

**Extended Data Figure 3.2:** Chronic chemogenetic activation of VMHvl$^{MC4R}$ neurons in intact females increases physical activity behavior during the inactive light period and reduces body weight.

**Extended Data Figure 4.1:** Increased physical activity and improvement in metabolic health markers in OVX VMHvl$^{MC4R::hM3Dq}$ females in response to acute and chronic CNO.

**Extended Data Figure 5.1:** Additional metabolic and expression data for conditional *Mc4r* rescue mice.

**Extended Data Figure 6.1:** Expression and physical activity levels in male and female CRISPRa$^{Mc4r}$ mice.

**Extended Data Table 1:** Brain region abbreviations

**Extended Data Table 2:** Statistical tests and results for main Figures 1-6

**Supplementary Video: CNO administered in the drinking water stimulates VMHvl$^{MC4R}$ neurons to increase physical activity.** Video recording of VMHvl$^{MC4R::hM3dq}$ (top) and VMHvl$^{Cre-}$ (bottom) female mice following addition of CNO-laden drinking water (0.25 mg/mL) during the inactive, lights-on period. Recordings have been sped up 20x.

**Krause et al., Extended Data 1.1**

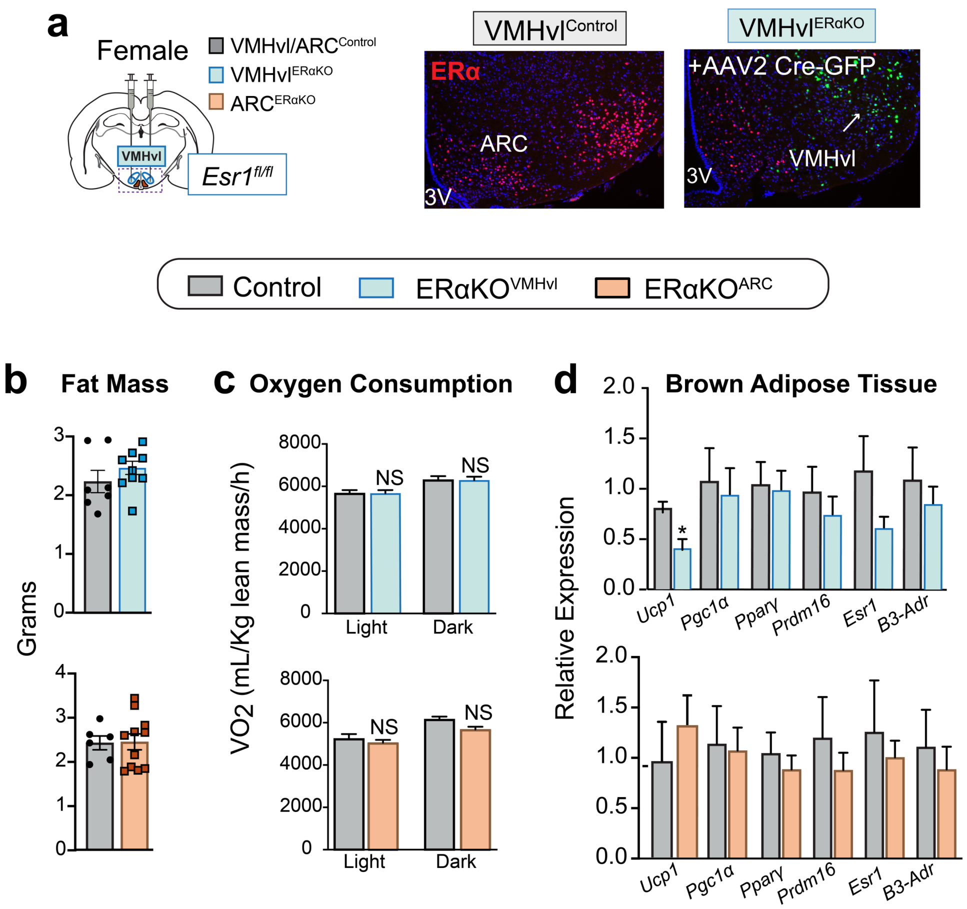


**Extended Data Figure 1.1: Stereotaxic-mediated VMHvl$^{ER\alpha KO}$ modestly affects interscapular brown adipose (iBAT) thermogenic gene expression in female mice maintained at ambient temperature. a**, Representative image of a successful hit confirmed post-mortem by loss of IF ERα expression (red) in the VMHvl (white arrow) with corresponding expression of GFP driven by the AAV2-Cre viral vector, as described in the Methods section. **b,** Equivalent fat mass and **c,** oxygen consumption rates in control, VMHvl$^{ER\alpha KO}$, and ARC$^{ER\alpha KO}$ female mice measured by EchoMRI and indirect calorimetry in CLAMS, respectively. **d**, Quantification of BAT thermogenic gene expression levels by qPCR in VMHvl$^{ER\alpha KO}$ and ARC$^{ER\alpha KO}$ female mice.

**Krause et al. - Extended Data 1.2**

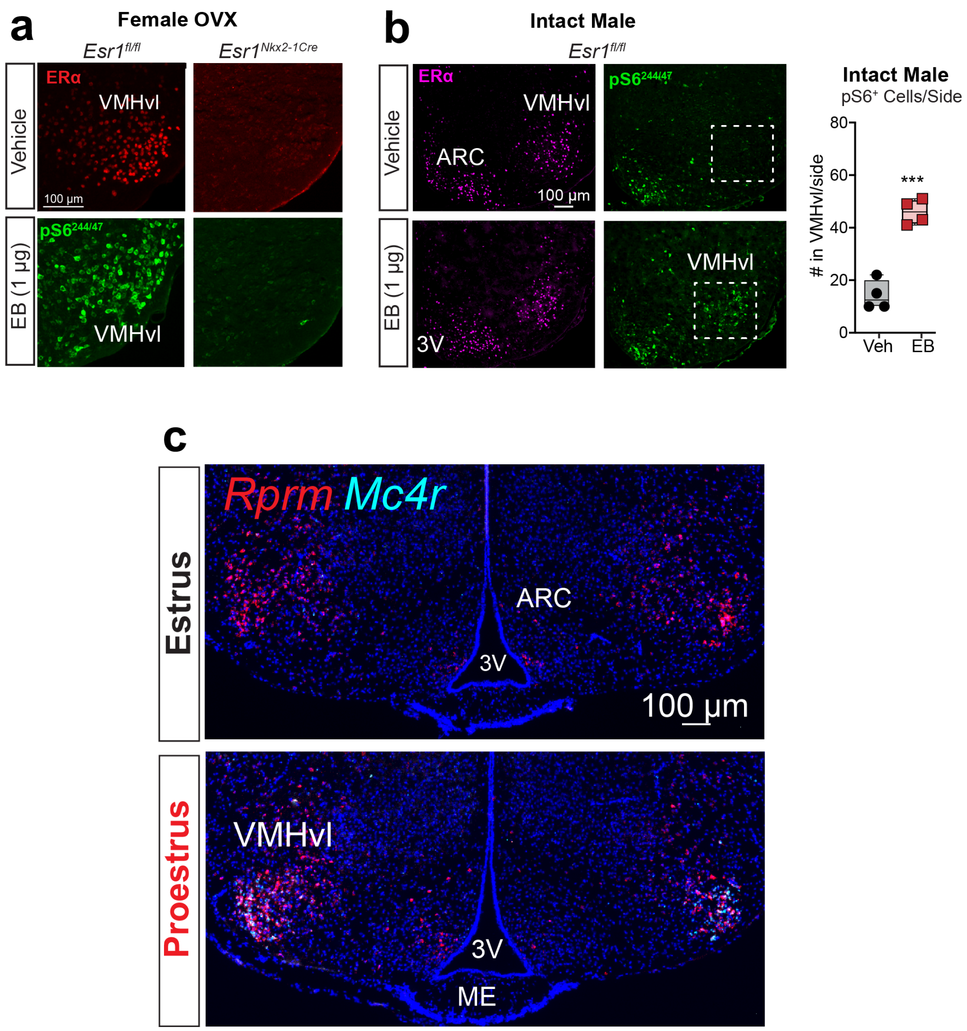


**Extended Data Figure 1.2: VMHvl pS6 induction depends on ERα and can be elicited in males by EB treatment. a,** IF of ERα (red) and pS6$^{244/47}$ (green) in the VMHvl of *Esr1$^{fl/fl}$* and conditional knockout (*Esr1$^{Nkx2\text{-}1Cre}$*) OVX female mice following 4 hrs post EB treatment. **b**, Images and bar graph showing increased number of pS6$^{244/47}$+ neurons in the VMHvl of male mice 4 hrs post EB treatment as done for OVX females (unpaired 2-tailed *t* test, $t_{(6)}$=8.569, $P$=0.0001). **c,** Full size images showing bilateral expression of *Mc4r* (blue) and *Rprm* (red) in intact females staged for estrus and proestrus. *Rprm* expression is unchanged in both stages of estrous.

**Krause et al., Extended Data 1.3**

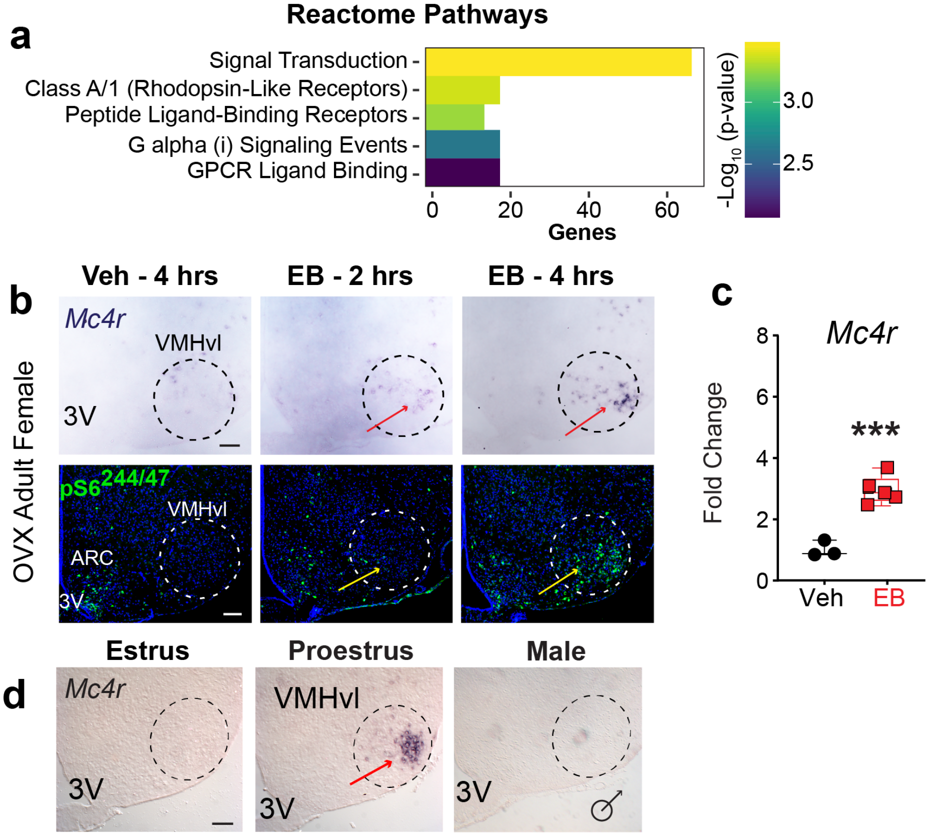


**Extended Data Figure 1.3: EB-dependent regulation of *Mc4r*. a**, Top five significantly enriched Reactome Pathways among DEGs in the VMHvl (vehicle vs. EB). **b**, Representative ISH (*Mc4r*, red arrows) and immunofluorescent (pS6, yellow arrows) staining in the VMHvl (dashed circle) from OVX female mice treated with vehicle for 4 hrs, EB for 2 hrs, or EB for 4 hrs. **c**, *Mc4r* expression levels in VMHvl from OVX females normalized to vehicle treatment (unpaired 2-tailed *t* test $t_{(6)}$=6.519, *P*=0.0006). **d,** ISH showing *Mc4r* expression in the VMHvl of an estrus female, a proestrus female, and an intact male.

**Krause et al., Extended Data 1.4**

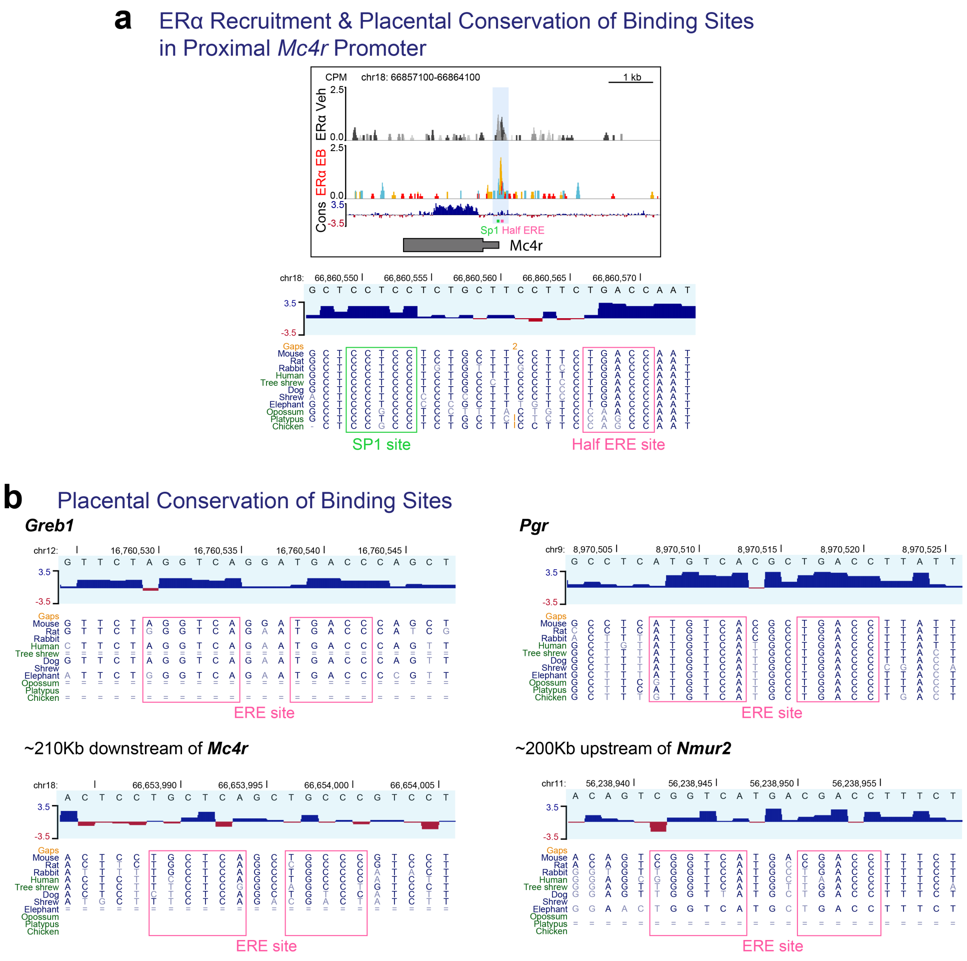


**Extended Data Figure 1.4: ERα binding sites in EB-sensitive target genes contain conserved ERE consensus sequences. a**, CUT&RUN CPM-normalized coverage track showing additional ERα binding site in the *Mc4r* promoter (1 of 3 replicates). **b**, Location and sequence conservation of half (*a*) and full (*b*) ERE consensus sites in target gene loci indicated by pink boxes. For all panels the genomic intervals containing ERE/SP1 sites are located within the ERα binding sites identified by CUT&RUN.

Krause et al. - Extended Data 2.1

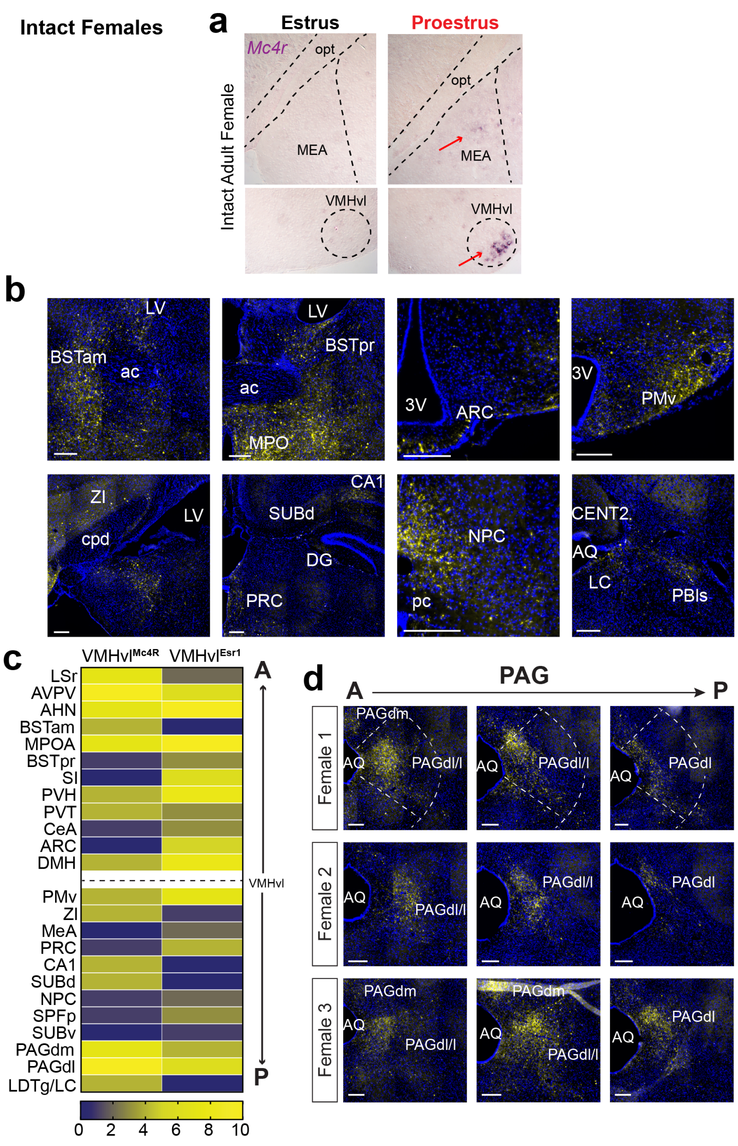


**Extended Data Figure 2.1: Sensitivity of *Mc4r* expression in the VMHvl and VMHvl$^{MC4R}$ neuron output. a**, Additional ISH comparing *Mc4r* induction in the VMHvl and MEA in estrus and proestrus females. **b**, Representative mYFP reporter expression in additional neuroanatomical regions. **c**, Heatmap from Fig. 3i rearranged to compare VMHvl$^{MC4R}$ and VMHvl$^{ER\alpha}$ projection intensity in target regions along the anterior-posterior axis. **d**, VMHvl$^{MC4R}$ projections to the PAG preferentially target the PAGdl/l and PAGdm.

**Krause et al. - Extended Data 3.1**

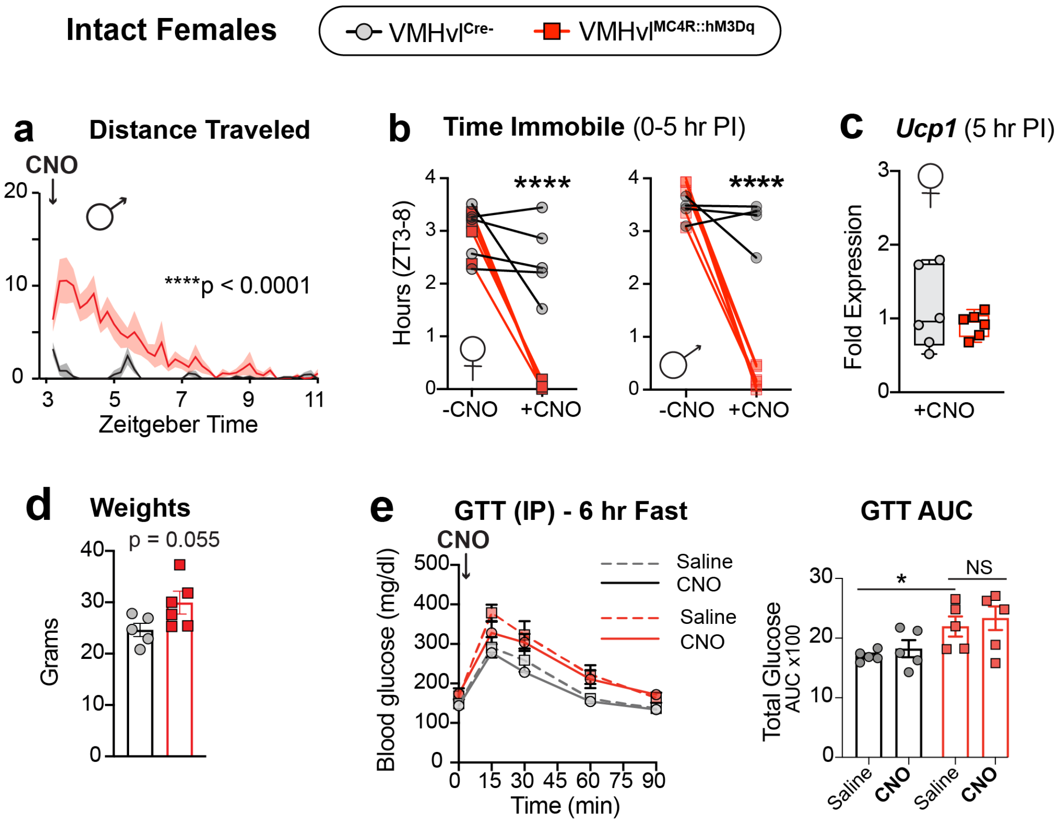


**Extended Data Figure 3.1: Chemogenetic activation of VMHvl$^{MC4R}$ neurons reduces sedentary behavior during the normally inactive light period but does not affect glucose homeostasis. a**, Distance traveled over time in male mice following a single injection of CNO (RM 2-way ANOVA interaction effect $F_{(39,312)}$=6.898, $P$<0.0001). **b**, Total time spent immobile in intact female and male VMHvl$^{Cre-}$ controls and VMHvl$^{MC4R::hM3Dq}$ mice (RM 2-way ANOVA female interaction effect $F_{(1,8)}$=33.89, $P$=0.0004, post hoc $P$<0.0001 and male interaction effect $F_{(1,8)}$=96.79, $P$=0.0005, post hoc $P$<0.0001). **c**, No differences were observed in *Ucp1* mRNA in the BAT from VMHvl$^{Cre-}$ and VMHvl$^{MC4R::hM3Dq}$ mice collected 1.5 hrs after a single CNO injection. **d**, Body weights for VMHvl$^{Cre-}$ and VMHvl$^{MC4R::hM3Dq}$ females at beginning of **e,** glucose tolerance testing with glucose levels shown over ninety-minute period for intact female cohorts treated with saline or CNO.

**Krause et al. - Extended Data 3.2**

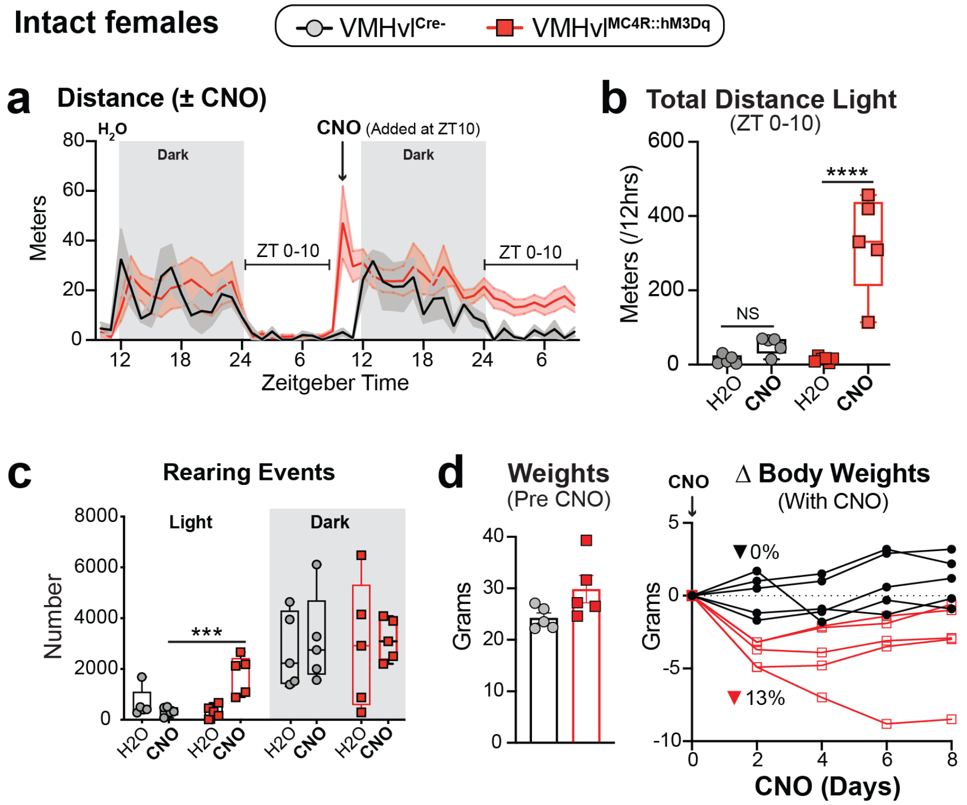


**Extended Data 3.2: Chronic chemogenetic activation of VMHvl$^{MC4R}$ neurons in intact females increases physical activity behavior during the inactive light period and reduces body weight. a,** Sustained physical activity increase across light/dark periods in VMHvl$^{MC4R::hM3Dq}$ females administered CNO-H2O as compared to VMHvl$^{Cre-}$ females or during exposure to plain drinking water ($H_2O$) **b,** Cumulative distance traveled and **c,** number of rearing events light/dark periods following administration of CNO or water during the light stage (b, RM 2-way ANOVA $F_{(1,8)}$=15.8, $P$=0.0041, post hoc $P$=0.0006 and c, RM 2-way ANOVA $F_{(1,8)}$=15.8, $P$=0.0041, post hoc $P$=0.0006). **d,** Starting body weights and weight change during continuous administration of CNO-$H_2O$ for intact females.

**Krause et al., Extended Data 4.1**

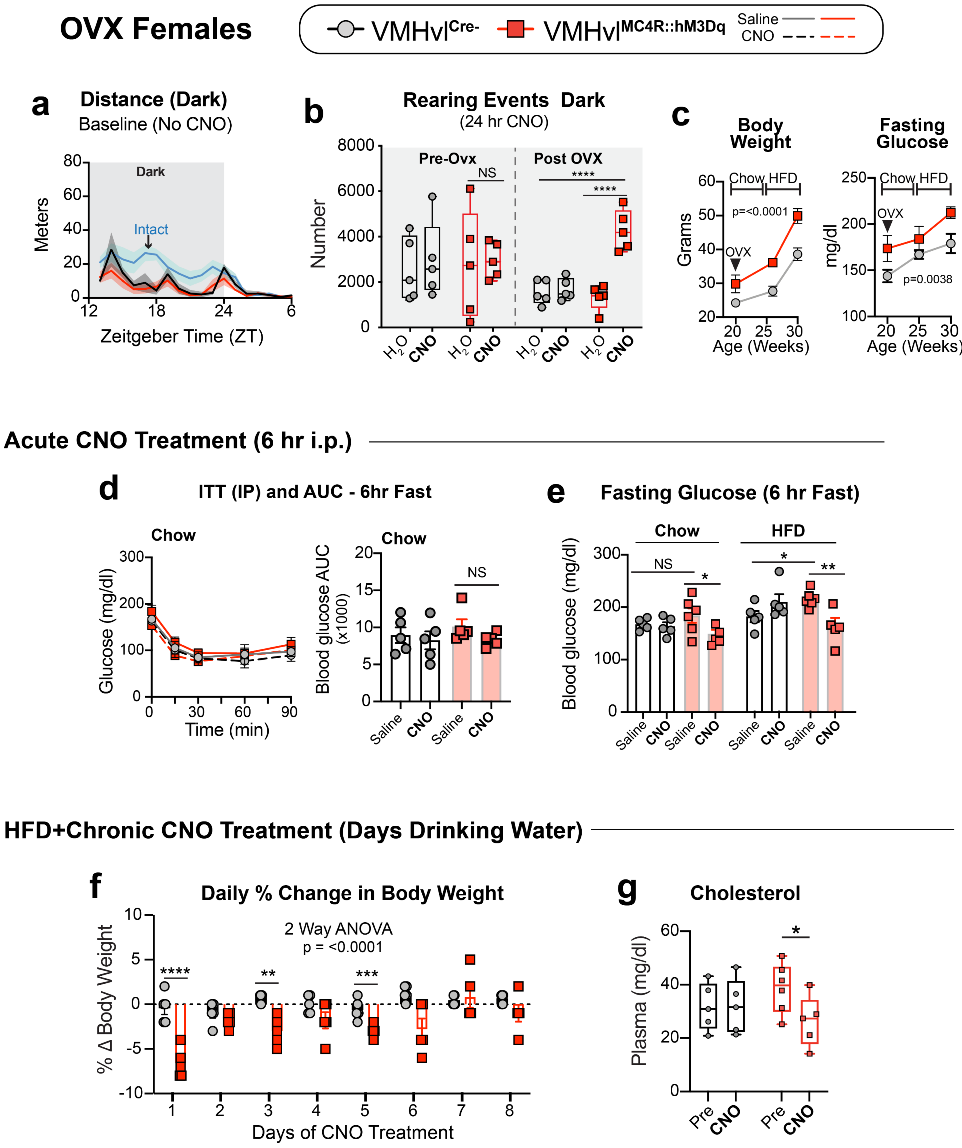


**Extended Data Figure 4.1: Increased physical activity and improvement in metabolic health markers in OVX VMHvl$^{MC4R::hM3Dq}$ females in response to acute and chronic CNO. a**, Distance traveled over time in OVX female VMHvl$^{Cre-}$ and VMHvl$^{MC4R::hM3Dq}$ mice during administration of plain H2O or CNO-H2O. **b,** Total dark period rearing events in intact and OVX females administered plain $H_2O$ or CNO-$H_2O$ drinking water (RM 2-way ANOVA interaction effect $F_{(1,8)}$=60.31, $P$<0.0001 post hoc $P$<0.0001). **c**, Body weights (RM 2-way ANOVA time effect $F_{(2,24)}$=49.51, $P$<0.0001; genotype effect $F_{(2,24)}$=33.50, $P$<0.0001) and fasting glucose levels (RM 2-way ANOVA time effect $F_{(2,26)}$=6.456, $P$=0.0053; genotype effect $F_{(1,26)}$=10.11, $P$=0.0038) in female mice after OVX and subsequent HFD feeding. **d,** Blood glucose and AUC during ITT in chow-fed OVX females following 6-hour fast and saline/CNO treatment. **e**, Blood glucose levels following a 6 hr fast in OVX females maintained on Chow/HFD following a single saline or CNO injection (RM ANOVA with mixed-effects model, note: one *Cre*$^{+}$ female with missed injection was excluded from CNO group, Chow: treatment effect $F_{(1,17)}$=5.038, *P*=0.0384, post hoc *P*=0.0179; and HFD: interaction effect $F_{(1,17)}$=20.47, *P*=0.0019, post hoc *P*=0.0073). **f**, Percent change in body weight in HFD-fed OVX females continuously administered CNO-laden drinking water (RM 2-way ANOVA interaction effect $F_{(7,64)}$=4.583, *P*=0.0003. **g**, Plasma cholesterol levels before (Pre) and after (Post) 8 days of continuous CNO-$H_2O$ exposure (RM ANOVA with mixed-effects model, note: one *Cre*$^{+}$ female with missed injection was excluded from CNO group, interaction effect $F_{(1,8)}$=5.502, *P*=0.0470, post hoc *P*=0.0203).

Krause et al., Extended Data 5.1

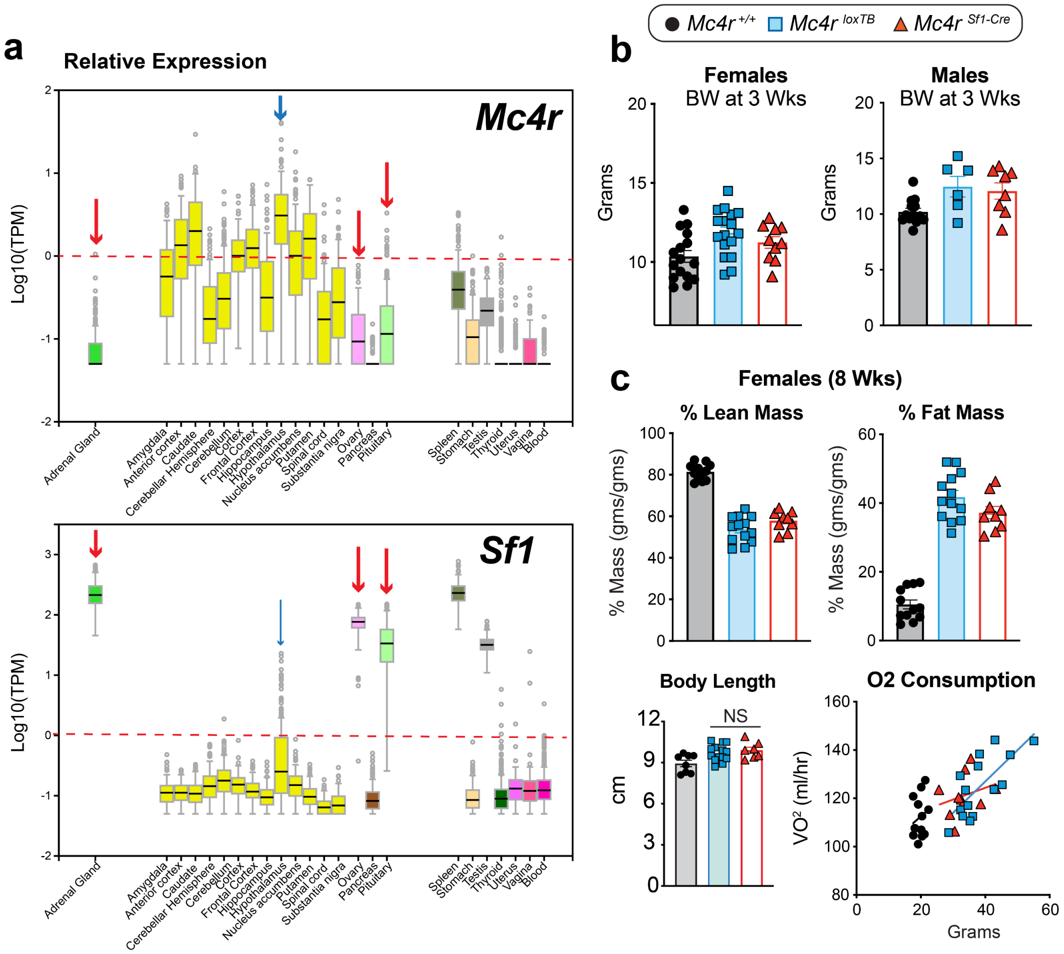


**Extended Data 5.1: Additional metabolic and expression data for conditional *Mc4r* rescue mice. a**, *Mc4r* and *Sf1* expression patterns from Genotype-Tissue Expression Project intersect specifically in the hypothalamus (blue arrows) and not in peripheral tissues (red arrows). **b**, Equivalent body weights within cohorts of female and male $Mc4r^{+/+}$, $Mc4r^{loxTB}$, and $Mc4r^{Sf1\text{-}Cre}$ mice at weaning. **c**, % lean (1-way ANOVA $F_{(2,31)}$=101.4, $P$<0.0001, post hoc: $Mc4r^{+/+}$ vs $Mc4r^{loxTB}$ $P$<0.0001, $Mc4r^{+/+}$ vs $Mc4r^{Sf1\text{-}Cre}$ $P$<0.0001, and $Mc4r^{Sf1\text{-}Cre}$ vs $Mc4r^{loxTB}$ $P$=0.0720) and % fat (1-way ANOVA $F_{(2,31)}$=104.2, $P$<0.0001, post hoc: $Mc4r^{+/+}$ vs $Mc4r^{loxTB}$ $P$<0.0001, $Mc4r^{+/+}$ vs $Mc4r^{Sf1\text{-}Cre}$ $P$<0.0001, and $Mc4r^{Sf1\text{-}Cre}$ vs $Mc4r^{loxTB}$ $P$=0.0769) body composition analysis (EchoMRI) in adult females of each genotype. Quantification of body length (1-way ANOVA $F_{(2,26)}$=5.465, $P$=0.0104, post hoc: $Mc4r^{+/+}$ vs $Mc4r^{loxTB}$ $P$=0.0196, $Mc4r^{+/+}$ vs $Mc4r^{Sf1\text{-}Cre}$ $P$=0.0171, and $Mc4r^{Sf1\text{-}Cre}$ vs $Mc4r^{loxTB}$ $P$=0.4904) in the three cohorts of adult female mice. Oxygen consumption (VO2) as a function of body weight in adult female mice.

**Krause et al., Extended Data 6.1**

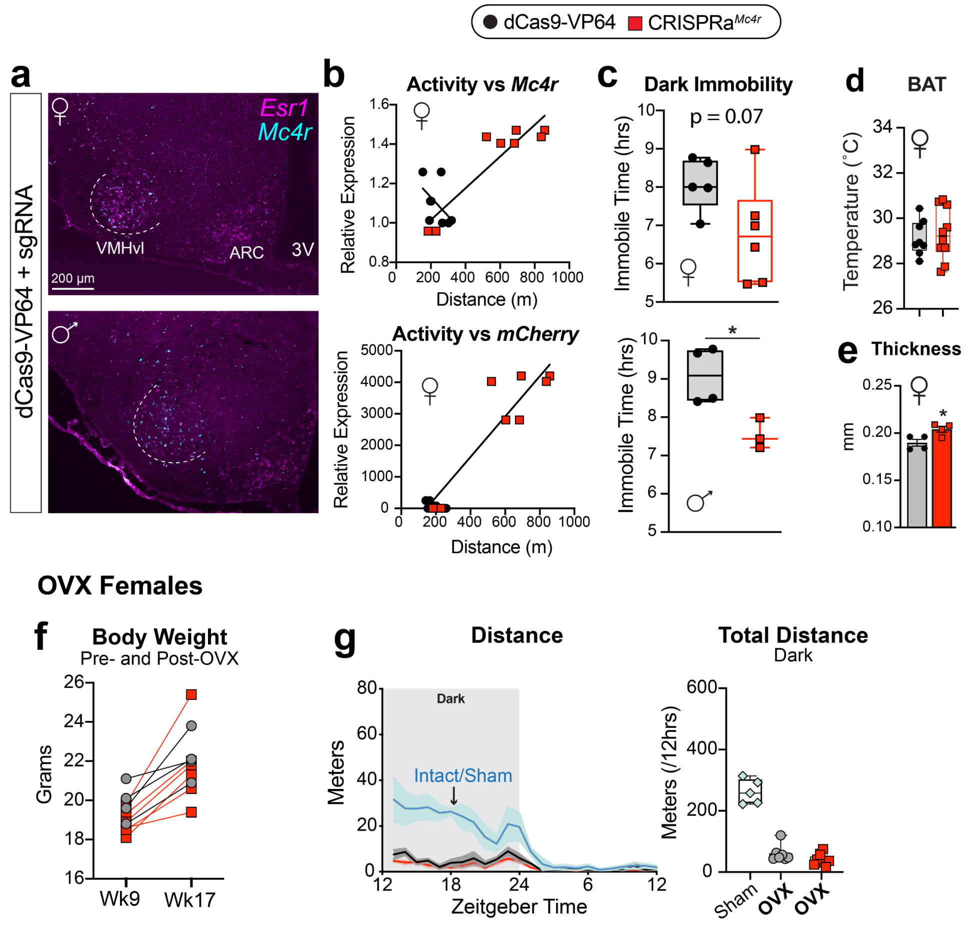


**Extended Data Figure 6.1: Expression and physical activity levels in male and female CRISPRa$^{Mc4r}$ mice.** **a**, Fluorescent ISH images from CRISPRa$^{Mc4r}$ female (top) and male (bottom) showing *Esr1* and *Mc4r* expression. Images are reproduced from Fig. 5b to show limited induction of *Mc4r* outside of the VMHvl target region. **b**, Dark phase (ZT12-24) physical activity levels (distance/12 hours) as a function of *Mc4r* or *mCherry* mRNA expression in microdissected VMHvl from control and CRISPRa$^{Mc4r}$ female mice. **c**, Time spent immobile during the 12 hour dark phase in control and CRISPRa$^{Mc4r}$ female (unpaired 2-tailed *t* test, $t_{(9)}$=2.015, $P$=0.0747) and male (unpaired 2-tailed *t* test, $t_{(5)}$=3.245, $P$=0.0228) mice. **d,** BAT surface temperatures in female control and CRISPRa$^{Mc4r}$ mice, repeated measurements at 30- and 60-min post-anesthesia. **e,** Cortical bone thickness for female cohorts (unpaired 2-tailed *t* Test, $t_{(6)}$=2.957, $P$=0.0254). **f**, Body weights in control and CRISPRa$^{Mc4r}$ females at wk 9 and at wk 17 after eight weeks of OVX. **g**, Distance traveled over 24 hours in OVX control and OVX CRISPRa$^{MC4R}$ compared to intact females (blue) with graph plotting total dark phase distance in intact, OVX control, and OVX CRISPRa$^{MC4R}$ females.

**Extended Data Table 1: Brain Region Abbreviations**

| Abbreviation | Full Name |
|---|---|
| 3V | Third Ventricle |
| 4V | Fourth Ventricle |
| ac | Anterior Commissure |
| AHN | Anterior Hypothalamic Nucleus |
| AQ | Cerebral Aqueduct |
| ARC | Arcuate Nucleus |
| AVPV | Anteroventral Periventricular Nucleus |
| B | Barrington’s Nucleus |
| BSTam | Bed Nuclei of the Stria Terminalis, anterior division, anteromedial area |
| BSTpr | Bed Nuclei of the Stria Terminalis, posterior division, principal nucleus |
| CA1d | Dosal Ammon's Horn, field CA1 |
| CeA | Central Amygdalar Nucleus |
| CENT2 | Central Lobule II |
| cpd | Cerebral Peduncle |
| d3V | Dorsal Third Ventricle |
| DG | Dentate Gyrus |
| DRN | Dorsal Raphe Nucleus |
| fr | Fasiculus Retroflexus |
| LC | Locus Coeruleus |
| LDTg | Laterodorsal Tegmental nucleus |
| LHb | Lateral Habenula |
| LSr | Lateral Septal Nucleus, rostral part |
| LV | Lateral Ventricle |
| ME | Median Eminence |
| MeA | Medial Amygdalar Nucleus |
| mlf | Medial Longitudinal Fascicle |
| MPO | Medial Preoptic Area |
| NPC | Nucleus of the Posterior Commissure |
| opt | Optic Tract |
| PAGdl/l | Periaqueductal Gray, dorsolateral/lateral |
| PAGdm | Periaqueductal Gray, dorsomedial |
| PAGvl | Periaqueductal Gray, ventrolateral |
| PBls | Parabrachial Nucleus, lateral division |
| pc | Posterior Commissure |
| PMv | Ventral Premammillary Nucleus |
| PVH | Paraventricular Hypothalamic Nucleus |
| PVT | Paraventricular Nucleus of the Thalamus |
| SUBd | Dorsal Subiculum |
| ZI | Zona Incerta |

### Extended Data Table 2: Statistical Tests and Results for Main Figures 1-6.

| Figure | Statistical Test | Result | Post Hoc Comparison(s) |
|---|---|---|---|
| **1a**, Body Weights | Unpaired 2-tailed *t* Test | $t_{(16)}$=2.365, *P*=0.0310 | |
| **1a**, X-Ambulatory | 2-Way ANOVA | interaction effect $F_{(1,15)}$=4.548, *P*=0.0499 | Dark Period, $VMHvl^{Control}$ vs $VMHvl^{ER\alpha KO}$ *P*=0.0014 |
| **1c**, VMHvl | Unpaired 2-tailed *t* Test | $t_{(7)}$=6.074, *P*=0.0005 | |
| **1d**, ARC | Unpaired 2-tailed *t* Test | $t_{(7)}$=1.562, *P*=0.1622 | |
| **1g**, *Mc4r* | 1-Way ANOVA | $F_{(2,14)}$=6.428, *P*=0.0105 | E vs P *P*=0.0163, P vs ♂ *P*=0.0189, and E vs ♂ *P*=0.7764 |
| **1g**, *Nmur2* | 1-Way ANOVA | $F_{(2,15)}$=8.469, *P*=0.0035 | E vs P *P*=0.0454, P vs ♂ *P*=0.0030, and E vs ♂ *P*=0.1438 |
| **1g**, *Esr1* | 1-Way ANOVA | $F_{(2,11)}$=10.18, *P*=0.0031 | E vs ♂ *P*=0.0033, P vs ♂ *P*=0.0374, and E vs P *P*=0.1650 |
| **2a**, VMHvl | Unpaired 2-tailed t Test | $t_{(16)}$=0.3669, *P*=0.7185 | |
| **2a**, MeA | Unpaired 2-tailed t Test | $t_{(6)}$=4.544, *P*=0.0039 | |
| **3b**, Female Distance/Time | RM 2-Way ANOVA | interaction effect $F_{(39,312)}$=11.96, *P*<0.0001 | |
| **3c**, Female Total Distance | RM 2-Way ANOVA | interaction effect $F_{(1,8)}$=27.48, *P*=0.0008 | CNO, $VMHvl^{Cre-}$ vs $VMHvl^{MC4R::hM3Dq}$ P<0.0001 |
| **3c**, Male Total Distance | RM 2-Way ANOVA | interaction effect $F_{(1,7)}$=36.27, *P*=0.0005 | CNO, $VMHvl^{Cre-}$ vs $VMHvl^{MC4R::hM3Dq}$ *P*<0.0001 |
| **3d**, Food Intake | RM 2-Way ANOVA | interaction effect $F_{(1,8)}$=3.502, *P*=0.0982 | CNO, $VMHvl^{Cre-}$ vs $VMHvl^{MC4R::hM3Dq}$ *P*=0.0489 |
| **3e** | RM 2-Way ANOVA | treatment effect | |
| **3f** | RM 2-Way ANOVA | interaction effect $F_{(1,8)}$=45.30, *P*=0.0001 | CNO, $VMHvl^{Cre-}$ vs $VMHvl^{MC4R::hM3Dq}$ *P*<0.0001 |
| **3h** | Unpaired 2-tailed t Test | $t_{(10)}$=6.555, *P*<0.0001 | |
| **3i** | Unpaired 2-tailed t Test | $t_{(10)}$=3.398, *P*=0.0068 | |
| **3j** | Unpaired 2-tailed t Test | $t_{(10)}$=2.727, *P*=0.0213 | |
| **4a**, Distance/Time | RM 2-Way ANOVA | interaction effect $F_{(11,88)}$=5.265, *P*<0.0001 | |
| **4b**, Total Distance | RM 2-Way ANOVA | $F_{(1,8)}$=73.6, *P*<0.0001 | Dark Period, $VMHvl^{Control}$ vs $VMHvl^{ER\alpha KO}$ *P*<0.0001 |
| **4b**, Total Distance | 1-Way ANOVA | $F_{(2,30)}$=27.88, *P*<0.0001 | intact vs OVX *P*<0.0001, intact vs OVX+CNO *P*=0.0023, and OVX vs OVX+CNO *P*<0.0001 |
| **4c**, Chow Diet | Unpaired 2-tailed *t* Test | $t_{(8)}$=9.122, *P*<0.0001 | |
| **4c**, HFD | Unpaired 2-tailed *t* Test | $t_{(8)}$=7.603, *P*<0.0001 | |
| **4d,** ITT Timecourse | RM 2-way ANOVA | interaction effect $F_{(12,68)}$=7.341, *P*<0.0001 | $VMHvl^{MC4R::hM3Dq}$ saline vs CNO, $T_{15}$ P=0.0009 and $T_{60}$ P=0.0318 |
| **4d**, ITT AUC | RM ANOVA with mixed effects model, 1 $Cre^{+}$ female with missed injection excluded from CNO group | interaction effect $F_{(1,8)}$=7.791, *P*=0.0235 | $VMHvl^{MC4R::hM3Dq}$ saline vs CNO *P*=0.0007 |
| **4e**, Weight Change Over Time | RM 2-Way ANOVA | interaction effect $F_{(8,64)}$=24.40, *P*<0.0001 | |

| **Figure** | **Statistical Test** | **Result** | **Post Hoc Comparison(s)** |
|---|---|---|---|
| **4e**, Weight Pre & Post CNO | RM 2-Way ANOVA | interaction effect $F_{(1,16)}$=4.781, $P$=0.0440 | VMHvl$^{MC4R::hM3Dq}$ Pre vs Post $P$=0.0169 |
| **4f** | RM ANOVA with mixed effect model, 1 Cre+ female with missed injection excluded from CNO group | interaction effect $F_{(1,17)}$=5.180, $P$=0.0361 | VMHvl$^{MC4R::hM3Dq}$ Pre vs Post $P$=0.0156 |
| **4g** | Nested *t* Test | $t_{(8)}$=3.447, $P$=0.0087 | |
| **5b** | 1-Way ANOVA | $F_{(2,9)}$=14.00, $P$=0.0017 | Veh+MT-II vs EB+MT-II $P$=0.0037, EB+Veh vs EB+MT-II $P$=0.0046 |
| **5d** | 1-Way ANOVA | $F_{(2,36)}$=25.25, $P$<0.0001 | *Mc4r*$^{+/+}$ vs *Mc4r*$^{loxTB}$ $P$<0.0001, *Mc4r*$^{+/+}$ vs *Mc4r*$^{Sf1\text{-}Cre}$ $P$<0.0001, and *Mc4r*$^{Sf1\text{-}Cre}$ vs *Mc4r*$^{loxTB}$ $P$=0.9851 |
| **5e**, 8 Weeks | 1-Way ANOVA | $F_{(2,41)}$=188.5, $P$<0.0001 | *Mc4r*$^{+/+}$ vs *Mc4r*$^{loxTB}$ $P$<0.0001, *Mc4r*$^{+/+}$ vs *Mc4r*$^{Sf1\text{-}Cre}$ $P$<0.0001, and *Mc4r*$^{Sf1\text{-}Cre}$ vs *Mc4r*$^{loxTB}$ $P$=0.0026 |
| **5e**, 13 Weeks | 1-Way ANOVA | $F_{(2,32)}$=226.6, $P$<0.0001 | *Mc4r*$^{+/+}$ vs *Mc4r*$^{loxTB}$ $P$<0.0001, *Mc4r*$^{+/+}$ vs *Mc4r*$^{Sf1\text{-}Cre}$ $P$<0.0001, and *Mc4r*$^{Sf1\text{-}Cre}$ vs *Mc4r*$^{loxTB}$ $P$=0.0029 |
| **5f** | 1-Way ANOVA | $F_{(2,25)}$=92.31, $P$<0.0001 | *Mc4r*$^{+/+}$ vs *Mc4r*$^{loxTB}$ $P$<0.0001, *Mc4r*$^{+/+}$ vs *Mc4r*$^{Sf1\text{-}Cre}$ $P$<0.0001, and *Mc4r*$^{Sf1\text{-}Cre}$ vs *Mc4r*$^{loxTB}$ $P$=0.1449 |
| **5g** | RM 2-Way ANOVA | interaction effect $F_{(2,30)}$=6.4, $P$=0.0047 | *Mc4r*$^{+/+}$ vs *Mc4r*$^{loxTB}$ $P$<0.0001, *Mc4r*$^{+/+}$ vs *Mc4r*$^{Sf1\text{-}Cre}$ $P$=0.0503 and *Mc4r*$^{Sf1\text{-}Cre}$ vs *Mc4r*$^{loxTB}$ $P$=0.0153 |
| **6c** | RM 2-Way ANOVA | interaction effect $F_{(1,30)}$=32.82, $P$<0.0001 | Dark Period, Control vs CRISPRa$^{Mc4r}$ $P$<0.0001 |
| **6d** | RM 2-way ANOVA | interaction effect $F_{(1,19)}$=16.51, $P$=0.0007 | Dark Period, Control vs CRISPRa$^{Mc4r}$ $P$<0.0001 |
| **6f** | Unpaired 2-tailed *t* Test | $t_{(11)}$=2.409, $P$=0.0347 | |
| **6g** | Unpaired 2-tailed *t* Test | $t_{(6)}$=3.498, $P$=0.0129 | |
